# Mean-field computational approach to HIV dynamics on a fitness landscape

**DOI:** 10.1101/518704

**Authors:** Hanrong Chen, Mehran Kardar

**Author notes:** Department of Physics and Astronomy, University of Pennsylvania, Philadelphia, PA, United States of America.

## Abstract

During infection by the human immunodeficiency virus (HIV), mutations accumulate in the intra-host viral population due to selection imposed by host T cell responses. The timescales at which HIV residues acquire mutations in a host range from days to years, correlating with their diversity in the global population of hosts, and with the relative strengths at which different regions of the HIV sequence are targeted by the host. In recent years, “fitness landscapes” of HIV proteins have been estimated from the global HIV sequence diversity, and stochastic simulations of *in silico* HIV infection, using these inferred landscapes, were shown to generate escape mutations whose locations and relative timescales correlate with those measured in patients with known T cell responses. These results suggest that the residue-specific fitness costs and epistatic interactions in the inferred landscapes encode useful information allowing for predictions of the dynamics of HIV mutations; however, currently available computational approaches to HIV dynamics that make use of realistic fitness landscapes are limited to these fixed-population-size stochastic simulations, which require many simulation runs and do not provide further insight as to why certain mutations tend to arise in a given host and for a given sequence background. In this paper, we introduce and examine an alternative approach, which we designate the evolutionary mean-field (EMF) method. EMF is an approximate high-recombination-rate model of HIV replication and mutation, in whose limit the dynamics of a large, diverse population of HIV sequences becomes computationally tractable. EMF takes as input the fitness landscape of an HIV protein, the locations and strengths of a host’s T cell responses, and the infecting HIV strain(s), and outputs a set of time-dependent “effective fitnesses” and frequencies of mutation at each HIV residue over time. Importantly, the effective fitnesses depend crucially on the fitness costs, epistatic interactions, and time-varying sequence background, thus automatically encoding how their combined effect influences the tendency for an HIV residue to mutate, in a time-dependent manner. As a proof of principle, we apply EMF to the dynamics of the p24 gag protein infecting a host whose T cell responses are known, and show how features of the fitness landscape, relative strengths of host T cell responses, and the sequence background impact the locations and time course of HIV escape mutations, which is consistent with previous work employing stochastic simulations. Furthermore, we show how features of longer-term HIV dynamics, specifically reversions, may be described in terms of these effective fitnesses, and also quantify the mean fitness and site entropy of the intra-host population over time. Finally, we introduce a stochastic population dynamics extension of EMF, where population size changes depend crucially on the fitness of strains existing in the population at each time, unlike prior stochastic simulation approaches with a fixed population size or a time-varying one that is externally defined. The EMF method offers an alternative framework for studying how genetic-level attributes of the virus and host immune response impact both the evolutionary and population dynamics of HIV, in a computationally tractable way.

**Author summary:** Fitness landscapes of HIV proteins have recently been inferred from HIV sequence diversity in the global population of hosts, and have been used in simulations of *in silico* HIV infection to predict the locations and relative timescales of mutations arising in hosts with known immune responses. However, computational approaches to HIV dynamics using realistic fitness landscapes are currently limited to these fixed-population-size stochastic simulations, which require many simulation runs and do not provide further insight as to why certain mutations tend to arise in a given host and for a given sequence background. Here, we introduce an alternative approach designated the evolutionary mean-field (EMF) method, which is an approximate high-recombination-rate model of HIV dynamics. It takes as input an HIV fitness landscape, the locations and strengths of a host’s immune responses, and the infecting HIV strain(s), and outputs a set of time-dependent “effective fitnesses” and frequencies of mutation at each HIV residue over time. We apply EMF on an example to show how features of the fitness landscape, relative strengths of host immune responses, and the HIV sequence background modify the effective fitnesses and hence the locations and time course of HIV mutations. We also develop a stochastic population dynamics extension of EMF where population size changes depend crucially on the fitness of strains existing in the population at each time. The EMF method enables more detailed study of how genetic-level attributes of the virus and host immune response shape the evolutionary and population dynamics of HIV, in a computationally tractable way.

## Introduction

During acute infection by the human immunodeficiency virus (HIV), the virus replicates rapidly and the viral load (concentration of HIV RNA particles in the blood plasma), a measure of the extent of HIV infection, rises exponentially to a peak 2–3 weeks post-infection [1]. Host cytotoxic T lymphocyte (CTL) responses are detectable just before this peak and expand as the viral load declines [1–3]. CTLs kill HIV-infected host cells through the recognition of HIV-derived peptides 8–10 amino acids long (called epitopes) that are presented on the surface of these cells bound to major histocompatibility complex (MHC) class I molecules [4]. While CTLs partially control HIV in an untreated host [5], they are unable to clear the infection: HIV mutants are generated stochastically via error-prone reverse transcription [6], and those with mutations within epitopes *escape* epitope-specific CTL recognition. Thus, CTLs impose selective pressures on the mutating intra-host HIV population and select for the emergence of escape mutations over time, which can be quantified by sequencing HIV in the blood of infected hosts [3, 7–9].

The timescales at which HIV escape mutations emerge in the intra-host population vary widely, ranging from days to years [1–3]. What is the origin of this variation, and how does it depend on properties of the virus and/or host immune response? Liu et al. [3] found a correlation between the rate of viral escape at an epitope, and the entropy of epitope sequences in the global population of hosts infected by the same group or subtype of HIV; they also found that the relative strength of CTL responses targeting each epitope (called the vertical immunodominance) plays an important role [3]. These results suggest that different HIV strains have different levels of replicative *fitness*: residues where mutations incur large fitness costs tend to be more conserved in the global population of hosts and also tend to acquire escape mutations more slowly; also, CTL responses impose additional fitness costs that are abrogated by escape mutations, and higher vertical immunodominance corresponds to larger costs imposed at an epitope. Using global diversity as an indicator for fitness is also supported by mutations toward the group or subtype consensus residue (called reversions), that tend to occur when HIV is transmitted to an MHC-mismatched host (who presents different epitopes from the donor) [10, 11] and continue over years of intra-patient evolution [9]. In recent years, residue-specific fitness costs of HIV mutants have been measured in cell culture [12–15] and from genetic variation within individual hosts [16], which were found to correlate with the global entropy of HIV residues belonging to the same group or subtype of HIV (although less conserved residues also tend to be more commonly targeted by CTLs [16], and low-fitness-cost mutants may sometimes be rarely observed globally [14, 15]). Apart from residue-specific fitness costs, epistatic interactions between residues were also shown to play an important role during HIV infection in particular for the Gag [17–21] and Nef [22] proteins, and this should be reflected in pairwise correlations in the global HIV sequence diversity. In recent years, “fitness landscapes” of various HIV proteins were inferred from this global diversity [12, 13, 23], and using such inferred landscapes, *in silico* simulations of HIV infecting patients with known CTL responses found a good correlation in the locations and relative timescales of escape mutations with those measured in the patients [24]. These results suggest that the fitness costs and epistatic interactions in the inferred landscapes encode useful information allowing for good predictions of the dynamics of HIV mutations given a host’s CTL responses, which may provide opportunities to inform the design of vaccine immunogens that promote combinations of escape mutations likely to be harmful for the virus [12, 21].

In this paper, we focus on methods for simulating HIV dynamics given a residue-specific fitness landscape such as those inferred previously [12, 13, 23]. Doing so is nontrivial as the presence of epistatic interactions deem it insufficient to simulate just one site or one epitope; the sequence background matters (as we will show later). Currently available approaches for simulating the dynamics of full HIV protein sequences including a fitness landscape are limited to the fixed-population-size stochastic simulations mentioned above [24]. However, a stochastic method requires performing many simulation runs (10^3^ in [24]) in order to start making predictions about the locations and timescales of HIV mutations, and it does not make clear *post hoc* why certain mutations tend to arise in a given host and for a given sequence background. Alternatively, compartmental models of HIV dynamics that consider multiple HIV strains and multiple CTL responses [25–28] could be extended to have one set of equations for each HIV strain in the population, but this would require a very large number of equations as the intra-host population expands and diversifies.

Here, we introduce and examine an alternative approach for simulating HIV dynamics given a fitness landscape, which we designate the evolutionary mean-field (EMF) method. The “mean-field” in EMF does not refer to chemical species reacting according to their mean concentrations (à la compartmental models), but to a class of approximations in statistical physics. Specifically, we map a model of HIV replication and mutation to a statistical physics model (following [29, 30]), a mean-field approximation of which gives a set of time-dependent “effective fields” or fitnesses, and estimates of the frequencies of mutation at each HIV residue over time. The EMF method enables efficient computation of the dynamics of mutations over time, for HIV proteins of realistic size and using realistic fitness landscapes, and unlike stochastic approaches (e.g. [24]) provide a single prediction for a given host and sequence background. Importantly, the effective fitnesses yielded by EMF depend crucially on the fitness costs, epistatic interactions, and time-varying sequence background, thus automatically encoding how their combined effect influence the tendency for an HIV residue to mutate. Analogous mean-field equations were derived in the high-recombination-rate limit (see e.g. Neher and Shraiman [31]), which is precisely the biological interpretation of the EMF approximation.

In the Results, we apply EMF as a proof of principle to study the dynamics of the p24 gag protein infecting a host whose CTL responses are known (from [3]). Specifically, we show how fitness costs and epistatic interactions in the fitness landscape, vertical immunodominance of CTL responses, and the sequence background modify the effective fitnesses and hence impact the locations and time course of HIV escape mutations.

These results are consistent with previous work employing stochastic simulations [24], and provide an alternative framework for understanding how genetic-level features of the virus and host CTL response combine to affect HIV dynamics. We also show how reversions and potential compensatory mutations may be described in terms of these effective fitnesses, and quantify the mean fitness and site entropy of the intra-host population over time. Since EMF is an approximate high-recombination-rate method, in S2 Appendix we perform fixed-population-size stochastic simulations with a finite recombination rate (as in [24]) to validate our results for this example.

Finally, a fixed-population-size simulation method (such as [24]) does not reflect that the viral load changes by several orders of magnitude during acute HIV infection [1], but computing population size changes by tracking the fitnesses of all HIV sequences in a large, diverse population is also computationally nontrivial. In the final part of this paper, we develop a stochastic population dynamics method based on the EMF approximation, where population size changes depend crucially on the fitness of strains existing in the population at each time, unlike prior stochastic simulation approaches with a fixed population size [24, 32] or a time-varying one that is externally defined [33]. Using this method on the example above, we show that the locations of HIV escape mutations are now stochastic, but the deterministic EMF method gives good predictions on average. This method also produces a characteristic exponential rise and fall of the HIV population size that is qualitatively consistent with the plasma viral load during acute infection. We end in the Discussion by discussing limitations, uses, and further extensions of EMF, including how to incorporate explicit target cell and CTL clone populations, like other compartmental models of HIV dynamics [25–28]. The EMF method offers an alternative framework for understanding how genetic-level features of HIV and the host CTL response impact HIV dynamics in terms of effective fitnesses, and enables a more detailed study of how the fitness landscape and sequence background impact the evolutionary and population dynamics of HIV, in a computationally tractable way.

## Methods

### The fitness landscape of an HIV protein

We model the fitness landscape of an HIV protein following prior work [12, 13, 24]: We describe HIV strains by their amino acid sequence, and in particular consider binary sequences *S* = (*s*_1_, …, *s*_*L*_) where *s*_*i*_ = 0 represents the consensus amino acid and *s*_*i*_ = 1 represents any other residue at position *i*, and *L* is the length of the protein. The binary approximation (used in [12] but not [13, 24]) is reasonable for highly conserved proteins such as p24 gag [12]; taking *s*_*i*_ to be one of four nucleotides or one of twenty amino acids is a straightforward extension, but beyond the scope of this paper.

We consider the intrinsic fitness of HIV sequence *S* to obey

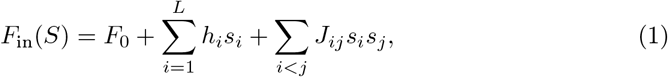

where *F*_0_ is the intrinsic fitness of the consensus sequence, {*h*_*i*_} are the fitness costs of mutations at site *i*, and {*J*_*ij*_} capture pairwise epistatic interactions between sites *i* and *j*^1^. We estimate the values of *F*_0_, {*h*_*i*_} and {*J*_*ij*_} in Parameter estimation.

During intra-host infection, the fitness of HIV sequence *S* is altered from its intrinsic value by host CTL pressure. Following [24], we model the effect of CTL-mediated killing of HIV-infected cells and HIV mutational escape by imposing a fitness cost *b*_*ε*_ on HIV sequences *S* with no mutation in epitope *ε*, which is abrogated if *S* contains at least one mutation within *ε*. Given a set of epitopes {*ε*} each beginning at site *i*_*ε*_ and of length *l*_*ε*_ amino acids, the within-host fitness of HIV sequence *S* is thus

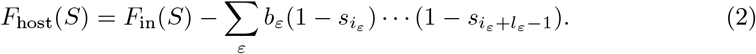

(This expression assumes that all targeted epitope sequences are (0, …, 0); if epitope *ε* instead has 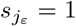, replace 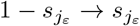.) In general, *b*_*ε*_ is time-dependent so *F*_host_(*S*) is time-dependent.

### Model of HIV dynamics on a fitness landscape

An HIV particle replicates by infecting a CD4^+^ host cell and reverse transcribing its RNA genome into DNA. HIV DNA is then integrated into host DNA and new copies of the virus are produced by the target cell. The ability of an HIV strain to replicate and infect new target cells in a host is described by its replicative fitness *F*_host_ (here modeled by Eq (2)). Also, reverse transcription is error-prone, and introduces random point mutations within the HIV genome with a probability that depends on the nucleotide-to-nucleotide substitution rates [6, 16].

Here, we model HIV replication and mutation by a discretized-time Markov model (see Fig 1). In one replication cycle (which takes roughly 1 day [34]), the effective number of offspring of HIV sequence *S* is *e*^*F*(*S*)^ where *F* = *F*_host_. Note that this is equivalent to solving the continuous-time equation 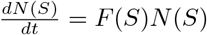 for unit time, where *N*(*S*) is the population size of sequence *S*; in the Discussion we elaborate on continuous-time extensions of our methods. Also note that we do not explicitly model infection of target cells or target cell dynamics; in the Discussion we discuss extensions that include explicit consideration of a target cell population.

To model error-prone reverse transcription, we consider that sequence *S* mutates with probability *μ*/site before replication (see Fig 1). Note that having mutations occur before replication is important for deriving the EMF equations below, but a model with replication before mutation can also be thought of as having one fewer round of mutations in the beginning, and hence both models should produce essentially the same results after many time steps if the mutation rate is small (not shown). Here we also consider *μ* to be independent of the residue, but we show in S1 Appendix that similar results are obtained when nucleotide-dependent substitution rates are accounted for.

First, let us write down the equations for replication and mutation of a single site *s*_*i*_. Suppose *s*_*i*_ = 0 has fitness *F*_0_ and *s*_*i*_ = 1 has fitness *F*_0_ + *h*_*i*_. Also suppose that *s*_*i*_ = 0 mutates to *s*_*i*_ = 1 or vice-versa with probability *μ*/generation. At time *k*, let the population size be 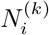 and the frequency of *s*_*i*_ = 1 in the population be 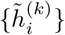. The population sizes of *s*_*i*_ = 0 and *s*_*i*_ = 1 at time *k* + 1 are thus given by

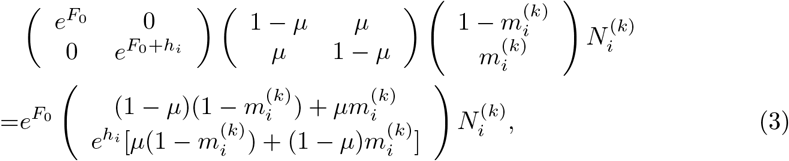

and their sum gives the total population size 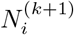 at time *k* + 1, with ratio

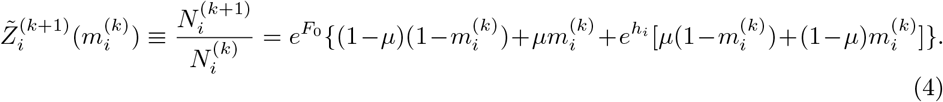

**Fig 1.**
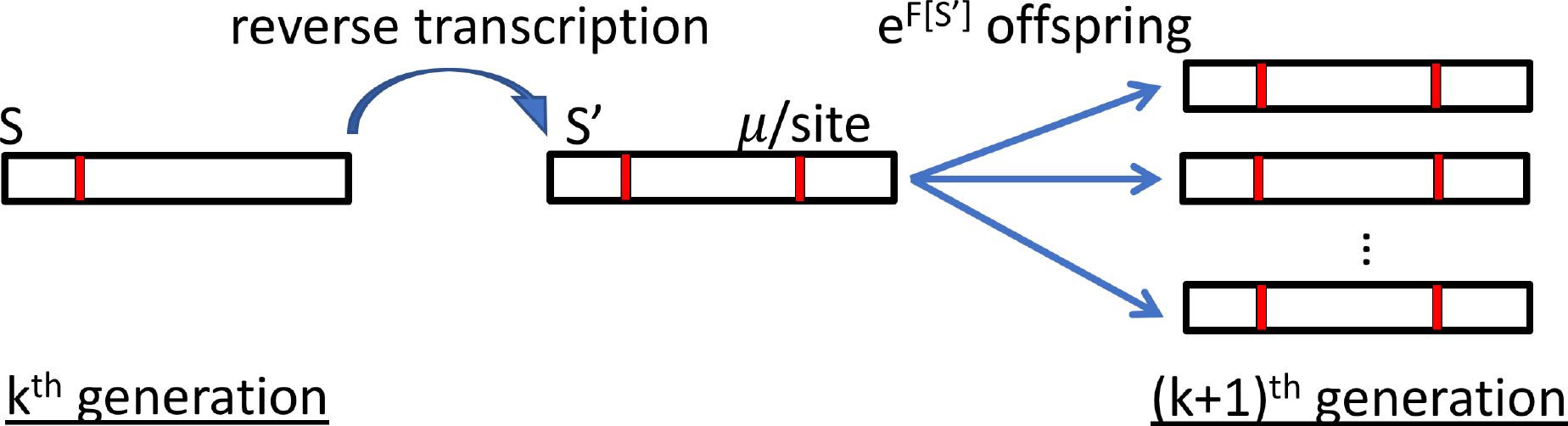
Model of HIV replication and mutation. In one replication cycle, HIV sequence *S* reverse transcribes its genome, introducing random point mutations with probability *μ*/site and producing sequence *S′*. Sequence *S′* produces *e*^*F*(*S′*)^ offspring in one generation. See Eq (5) for the transition matrix describing this process.

Thus, for HIV sequences of length *L* replicating according to a fitness landscape Eq (2), the number of offspring with sequence *S*^(*k*+1)^ produced by a single sequence *S*^(*k*)^ in 161 one generation (or equivalently, the ratio between their populations) is given by

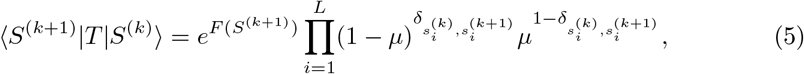

where 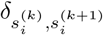 is the Kronecker delta. The 2^*L*^ × 2^*L*^ transition matrix *T* is not a transition probability matrix (which would require ∑_*S′*_〈S′|T|S〉 = 1), but the transition matrix of a model with *time-varying* population size. The state at each time is a vector 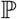 of populations of the 2^*L*^ sequences *S*.

Now, suppose we know the fitness landscape *F*_host_ and the initial population 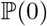. Our goal is to characterize 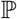 at future times, specifically to answer two kinds of questions:

1. What is the steady state at long times? Which escape mutations arise? Are there reversions or compensatory mutations?
2. What are the dynamics towards the steady state? What is the order and timescale of mutations? Are there transient dynamics?

The most direct method of solution is to find 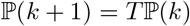 for *k* = 0, …, *n*, from which we can compute, for example, the statistics of mutations at site *i* over time. This involves, at time *k*, finding the marginal frequency

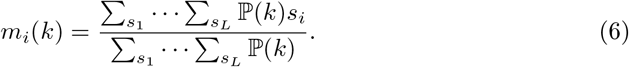

Equipped with this information, we can for example predict the relative probabilities of making each escape mutation in an epitope.

However, this direct method is computationally intractable. For an HIV protein of length *L* = 200, there are 2^200^ ≈ 10^60^ possible sequences, so brute-force matrix multiplication, matrix decompositions of *T*, or computation of Eq (6) by direct summation aren’t feasible in general. In practice, if the population size is not prohibitively large or if 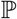 is sparse, then all sequences existing in the population may be tracked individually. In particular, if *μ≪N*1 (where 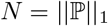 is the population size), successive selective sweeps occur and the population may be described by a dominant strain at each time [35]. However, this is not the regime HIV is in during intra-host infection, when the population continually accumulates diversity through mutations that are selected upon by strong CTL pressure. A commonly used method when *μN* ≳ 1 is to perform stochastic simulations with a fixed population size *N*. In S2 Appendix, we perform such (Wright–Fisher) simulations following the methods of [24], for the example considered in Results.

### The evolutionary mean-field (EMF) method

In the following, we solve 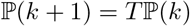 for *k* = 0, …, *n* approximately, which directly yields estimates of Eq (6) for all HIV residues over time.

To do this, we first map the above model of HIV replication and mutation to a statistical physics model (first introduced in [29, 30]). Using the identity 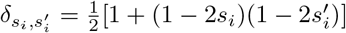, we rewrite Eq (5) in the form 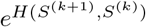, with energy (Hamiltonian) given by

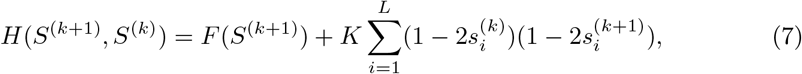

where 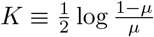, and we have omitted a constant term 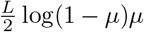 for notational clarity. The total number of offspring produced by sequence *S*^(*k*)^ in one generation is given by

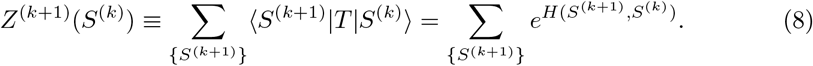

Note that *H*(*S*^(*k*+1)^, *S*^(*k*)^) depends on both *S*^(*k*)^ and *S*^(*k*+1)^, while the ‘partition function’ *Z*^(*k*+1)^ has the 2^*L*^ possible sequences {*S*^(*k*+1)^} summed over. Extending to multiple generations, the Hamiltonian has a structure shown in Fig 2, left, with sites (spins) within each generation interacting according to *F*(*S*), and spins along the time dimension having nearest-neighbor couplings *K* [29, 30]. The partition function (summing over {*S*^(1)^}, …, {*S*^(*n*)^}) is then the total number of offspring produced by a single sequence *S*^(0)^ over *n* generations.

Now, consider an approximate Hamiltonian

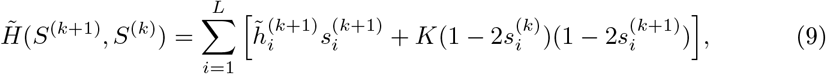

where the “effective fields” or fitnesses 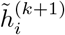 at sites *i* = 1, …, *L* are to be determined. Unlike Eq (7), here the spins *i* are noninteracting (see Fig 2, right). This simplifies computation of the total number of offspring produced by sequence *S*^(*k*)^ in one generation, to

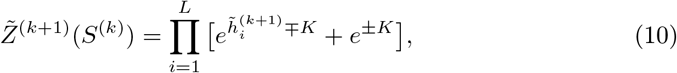

which is now a product of *L* terms instead of a sum over 2^*L*^ terms (cf. Eq (8)). In Eq (10), the upper sign is for 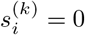 and the lower sign is for 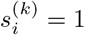.

**Fig 2.**
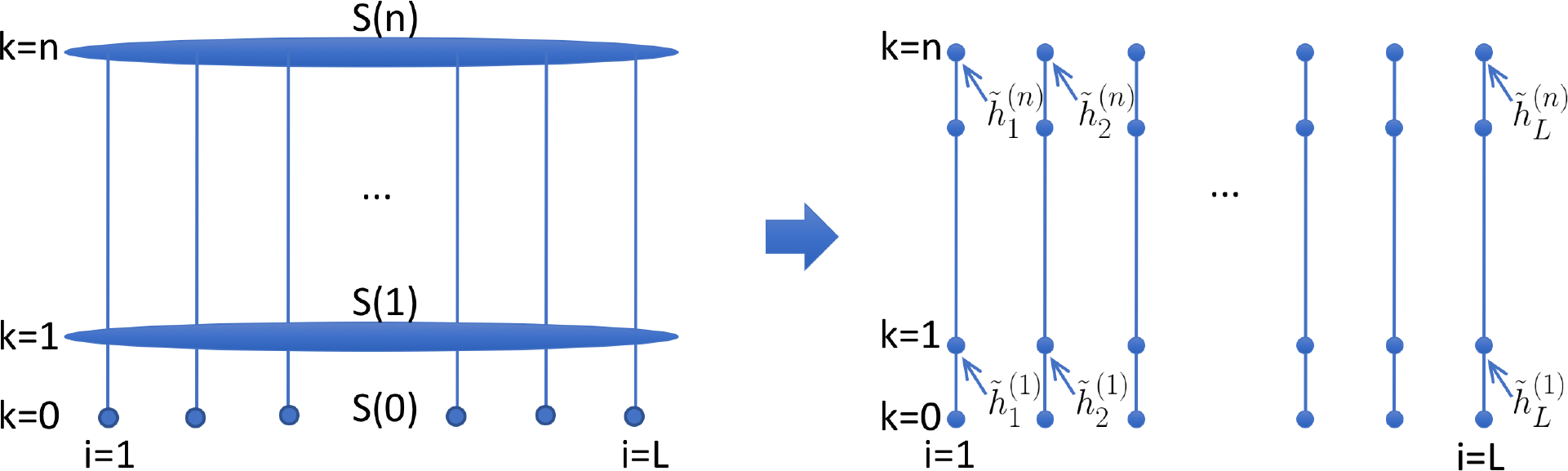
The evolutionary mean-field approximation. Left: A statistical physics model with Hamiltonian *H*(*S*^(*n*)^, …, *S*^(0)^), generalizing Eq (7) to multiple generations. Right: A schematic of the evolutionary mean-field (EMF) approximation. The “effective fitnesses” 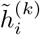 depend on the time-varying sequence background in a nontrivial manner (see Eqs (14) and (16)).

We want to define effective fitnesses 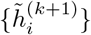 such that *Z*(*S*^(*k*)^) and 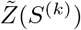 are as “close” as possible. To do this, we make use of Gibbs’ inequality [36], which states that the Kullback–Leibler divergence of two distributions *P* and 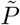 is nonnegative, i.e. 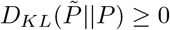. Using the forms *P*(*·*) = *e*^*H*(·)^/*Z* and 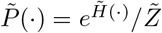 Gibbs’ inequality becomes

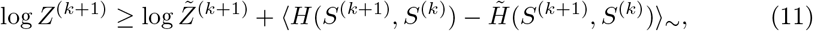

Where 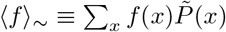 is the expectation value of *f* with respect to the approximate model. (We note here that Shekhar et al. [37] used Eq (11) in similar work to study the relation between the prevalence and intrinsic fitness landscapes of HIV; they used a different 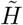 involving *F*_in_ instead of Eq (9), and their resulting equations do not reduce the computational complexity of the solution, which is our goal here, because their 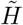 is still interacting.)

For the within-host fitness landscape Eq (2),

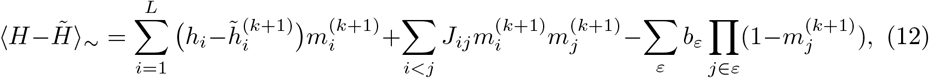

where we have defined 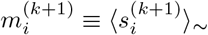 and 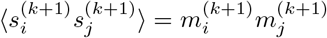 because sites are decoupled in the approximate model. Thus, extremizing the RHS of Eq (11) w.r.t. 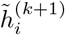, and using Eqs (10) and (12), leads to

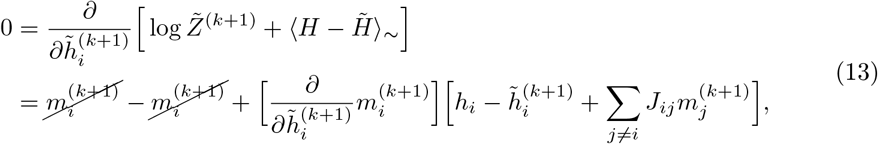

for sites *i* outside of epitopes giving

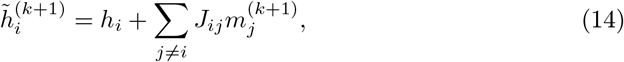

and

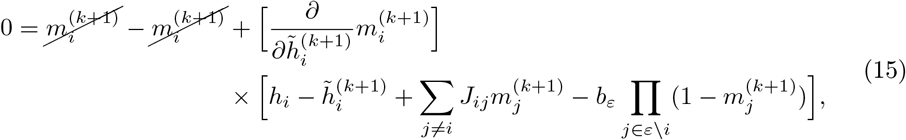

for sites *i* in epitope *ε* giving

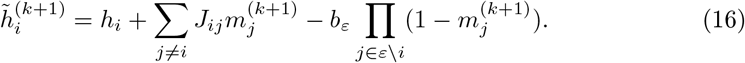

Equation (16) again assumes that the targeted epitope sequence is (0, …, 0); replace 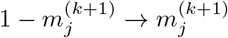 if *ε* contains a site *j* with *s*_*j*_ = 1.

The 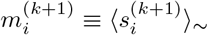 are the frequencies of mutation at site *i* and time *k* + 1 in the approximate model. For 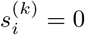, we find by differentiating Eq (10) w.r.t. 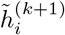 that

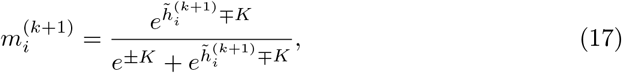

where again the upper sign is for 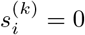 and the lower sign is for 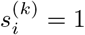 For generic 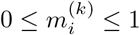, we obtain

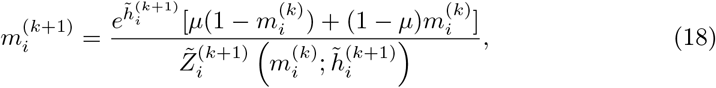

which is analogous to the one-site case (see Eqs (3) and (4)).

To summarize, the EMF method starts with the initial frequencies of HIV mutations 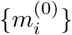, and computes the *L* effective fitnesses 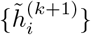 (Eqs (14) and (16)) and frequencies of mutations 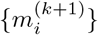 (Eq (18)) at times *k* + 1 recursively from the 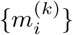 at the previous time *k*. While computing the marginal frequencies at each time by Eq (6) involves summing over 2^*L*^ terms, which is intractable for HIV sequences of realistic size, Eq (18) contains a sum of no more than two terms in the denominator, however requiring simultaneous solution with Eqs (14) and (16) (because 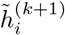 depends on 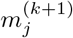 for *j /*= *i* which in turn depend on 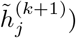). A dynamic-programming-like method of solution is to iterate between 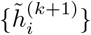 and 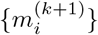 until convergence for each *k* (not shown). However, if the 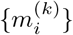 do not change drastically with time, a computationally more efficient implementation is to replace 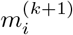 with 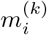 in Eqs (14) and (16), and to “shoot” forward in time *k*, iterating between 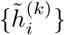 and 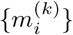 for larger and larger *k*:

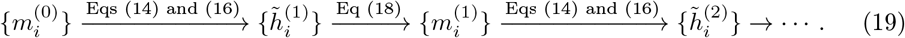

### EMF population dynamics method

Finally, the EMF method also allows for the computation of the *population dynamics*, since *Z*^(*k*+1)^ (Eq (8)) is the ratio of total population sizes between times *k* + 1 and *k*. While computing *Z*^(*k*+1)^ involves summing over 2^*L*^ terms, the RHS of Eq (11) gives a computationally tractable lower bound:

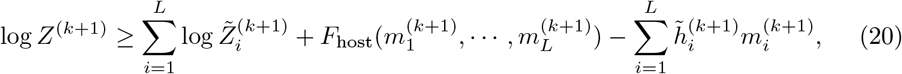

where 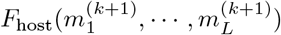 is the mean fitness of the population at time *k* + 1 within the EMF approximation.

Motivated by prior stochastic simulation approaches [24, 32, 33], we thus define a *stochastic* population dynamics method based on EMF by taking the population size at generation *k* + 1, *N*^(*k*+1)^, to be Poisson-distributed (following e.g. [33]) with mean *N*^(*k*)^*Z*^(*k*+1)^. Importantly, population size changes depend crucially on which mutations exist and arise in the population at each time. We also draw the frequencies of mutations 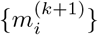 at time *k* + 1 from a binomial distribution (following e.g. [24, 32, 33]) with *N* ^(*k*+1)^ trials and probabilities given by Eq (18).

### Interpretation of EMF as a high-recombination-rate model of HIV dynamics

The EMF method solves the model of HIV replication and mutation defined by Eq (5) approximately by a model with *L independently* evolving sites experiencing *time-dependent* fitnesses 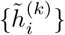, yielding estimates of the frequencies of mutation 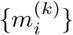 at all HIV residues over time. While this approximation was motivated by statistical physics, mean-field equations analogous to Eq (14) were derived for genotype dynamics in the high-recombination-rate limit (see Neher and Shraiman [31]). Indeed, in this limit the probability that mutations at sites *i* and *j* are jointly observed on a sequence is simply equal to the product of their frequencies, *m*_*i*_*m*_*j*_ (called linkage equilibrium [38]). Because EMF approximates sequence dynamics by the dynamics of independently evolving sites, it should be viewed as a high-recombination-rate model of HIV dynamics.

HIV does undergo recombination during intra-host infection: it switches RNA templates during reverse transcription at a rate estimated at 2.8 events/genome/cycle [39], and this may generate novel recombinant sequences if a target cell is coinfected by multiple HIV strains [40]. The effective HIV recombination rate was estimated from intra-host genetic variation to be 1.4 × 10^−5^/nucleotide/day [41], which is of the same order as its mutation rate. Recombination plays an important role during HIV infection because separate escape mutations recombining onto the same genome generate fitter viruses that escape multiple epitopes [42], and so models of HIV dynamics with high recombination rates would lead to higher escape rates from multiple epitopes [32]. Because EMF is a high-recombination-rate approximation, escape mutations predicted by this method should occur on a faster timescale than in models with a finite recombination rate (e.g. [24, 32]), and so the transient dynamics produced by EMF may be inaccurate, but the locations and relative timescales of mutations caused by fitness effects should nevertheless be comparable and consistent. Indeed, in S2 Appendix we demonstrate consistency of the results presented for the example below with those obtained using the stochastic simulation method of [24] with a finite recombination rate.

### Parameter estimation

#### Prevalence landscape of the p24 gag protein

p24 gag, which encodes the HIV capsid, is a highly conserved protein, for which binarized amino acid sequences is a reasonable approximation [12]. We follow the methods of [24] to infer the prevalence landscape of p24 from an alignment of HIV-1 group M subtype B protein sequences (downloaded from [43]). To improve data quality, all sequences with >5% gaps or ambiguous amino acids were excluded, and all remaining ambiguous amino acids were imputed by the consensus at that position [24]. The sequences were then binarized such that the most common (consensus) residue at each position was relabeled 0, and any mutation (or gap) was relabeled 1. To prevent multiple sequences drawn from the same patient from introducing biases in the sample mean and second moment, we weighted each sequence by one divided by the number of samples from that patient in the alignment [24]. We describe the distribution of p24 sequences by 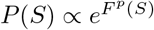 where the prevalence landscape obeys the form

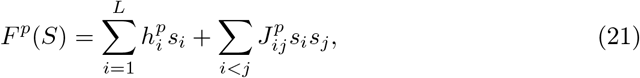

which is the maximum-entropy distribution given the empirical means and second moments of mutation (see [12, 13]). We inferred the values of 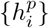 and 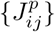 using available techniques [44, 45], and the resulting prevalence landscape is shown in Fig 3.

**Fig 3.**
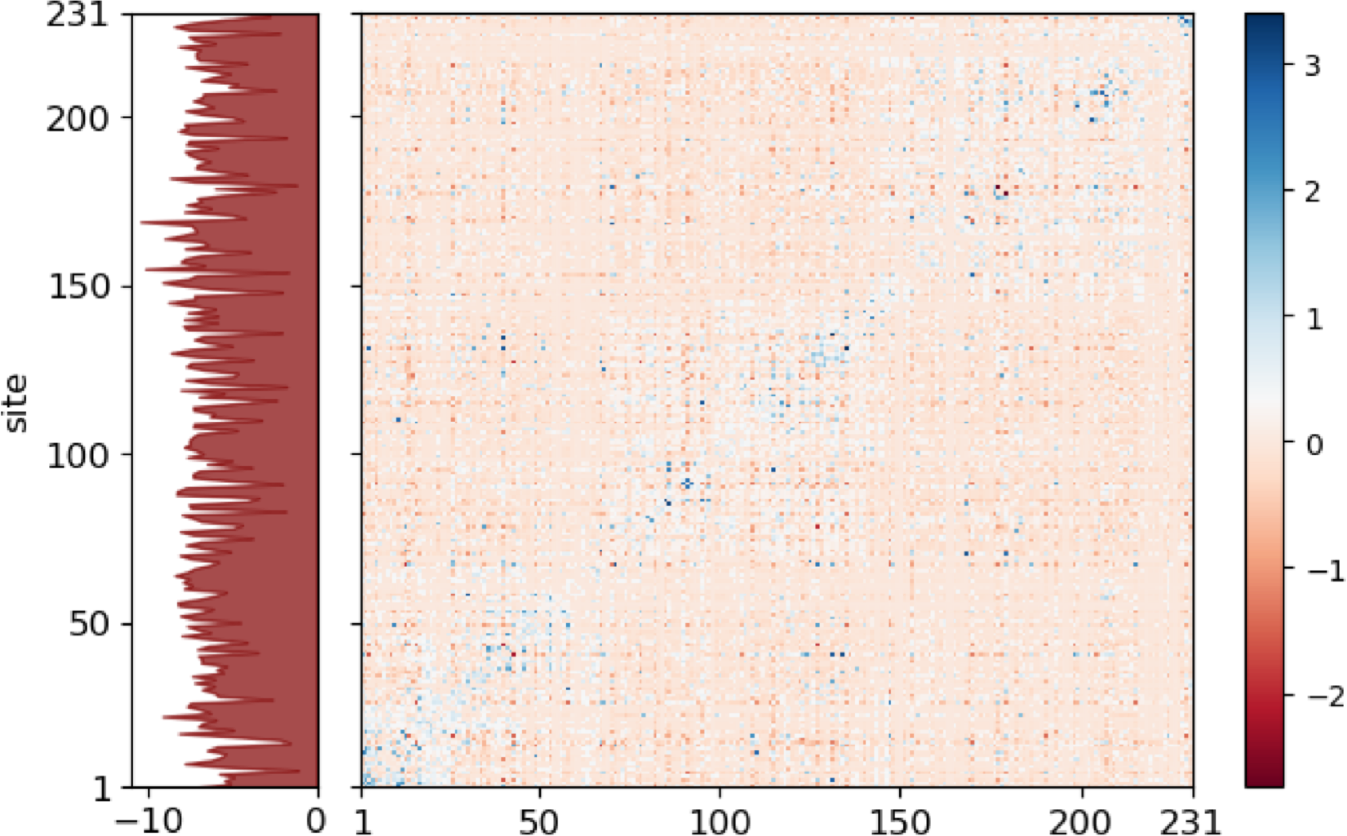
The prevalence landscape of p24. Left panel: Fields 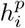 along the p24 sequence (length *L* = 231). All 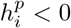 because p24 is highly conserved so all *s*_*i*_ = 1 are observed less frequently than *s*_*i*_ = 0. Right panel: Color map of couplings 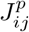; positive couplings are in blue and negative couplings are in red.

#### Estimating the proportionality factor between prevalence and fitness landscapes,*β*, and fitness of consensus,*F*_0_, from replicative capacity measurements

Shekhar et al. [37] showed via *in silico* simulations and a variational argument that the prevalence landscape (Eq (21)) and intrinsic fitness landscape (Eq (1)) of HIV are proportionally related in certain regimes, particularly when the global population of hosts mounts very diverse immune responses (which is to some extent satisfied; see e.g. [16]). We thus relate the 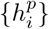 and 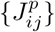 of the prevalence landscape and the {*h*_*i*_}. and {*J*_*ij*_} of the fitness landscape by a proportionality factor *β*, i.e. 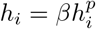 and 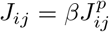 Note that *β* has units of inverse time, because 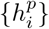 and 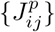 are dimensionless while {*h*_*i*_} and {*J*_*ij*_} describe replicative fitness costs and epistatic effects with units of inverse time.

To estimate *β*, we first turned to Mann et al. [13], who found a correlation between the measured replicative capacities *RC*(*S*) of a number of p24 mutants *S* in cell culture, and their prevalence landscape values *F*^*p*^(*S*). Specifically, they found a proportionality factor of 0.07 between *RC*(*S*)/*RC*(*S*_NL4-3_) and *F*^*p*^(*S*) − *F*^*p*^(*S*_NL4-3_), where NL4-3 is a reference strain with mutations at sites 120 and 208 in p24 with respect to the HIV-1 group M subtype B consensus sequence. Given that *RC*(*S*_NL4-3_) = 1.5 day^−1^ [13], we obtain *β* = 0.07 × 1.5 = 0.11 day^−1^. We found that this value of *β* produces relative fitness costs (*F*_0_ − *F*_in_(*S*))*/F*_0_ distributed around 40–50%, which are significantly larger than those inferred from intra-host variation in another study that were (broadly) distributed around 10% [16]. Because replicative capacity measurements in cell culture may not quantitatively equal viral growth rates in a host, motivated by [16] we instead took *β* = 0.023 day^−1^, which gives relative fitness costs of all single and double mutants shown in Fig 4.

Using *F*^*p*^(*S*_NL4-3_) = −4.13, *RC*(*S*_NL4-3_) = 1.5 day^−1^, and *β* = 0.023 day^−1^, we solve *F*_0_ − 1.5 = *β*(0 − *F*^*p*^(*S*_NL4-3_)) to find *F*_0_ = 1.6 day^−1^. We performed the following back-of-the-envelope check: around the time of peak viremia—roughly 2 weeks after infection—there are ~10^10^ virus particles in a host (because there is a peak viral load of ~10^6^ RNA copies/ml of blood [1] and ~5 liters of blood in a human). Solving exp(*F* × 14 days) = 10^10^ gives *F* ≈ 1.6 day^−1^, which is consistent.

**Fig 4.**
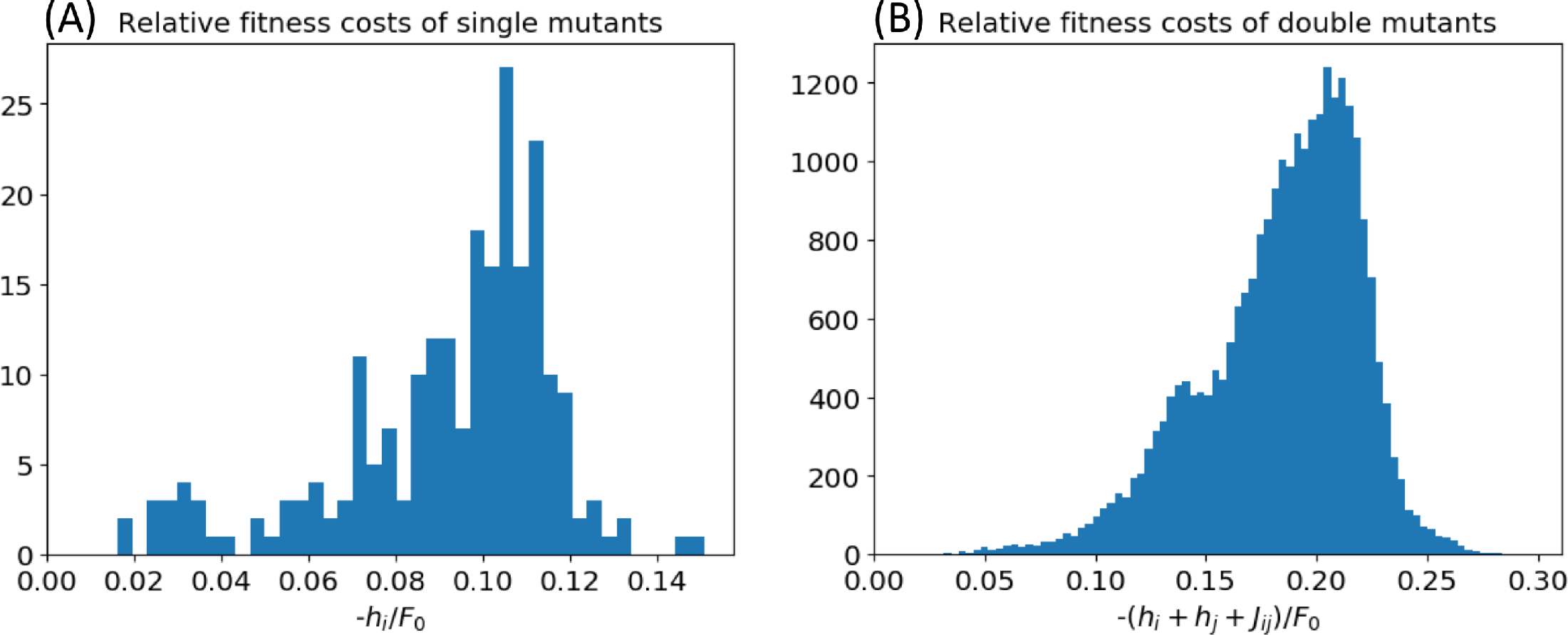
Histograms of relative fitness costs of all single and double mutants according to the fitness landscape Eq (1), using the prevalence landscape of Fig 3, *β* = 0.023 day^−1^, and *F*_0_ = 1.6 day^−1^.

#### The CTL response, *b*_*ε*_(*t*)

For the kinetics of the CTL response at epitope *ε*, *b*_*ε*_(*t*), we assumed the following Hill-like functional form^2^:

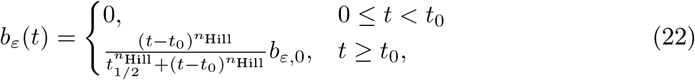

where the CTL response is activated *t*_0_ days post-infection, increases to a maximum of *b*_*ε*,0_ at long times, and *t*_1/2_ and *n*_Hill_ parametrize the rate of this increase. As a systemic HIV infection begins around 5–10 days post-infection and CTL responses emerge 2–3 weeks post-infection [1], we took *t*_0_ = 7 days, *t*_1/2_ = 7 days, and *n*_Hill_ = 2.

For the overall magnitude of the CTL response *b*_tot_ = ∑_*ε*_ *b*_*ε*,0_, we found that a range of values lead to realistic-looking population dynamics curves during the first weeks of infection (see S1 Fig). We believe this is biologically plausible as different untreated hosts presumably have variations in the magnitudes and timescales of their CTL responses, yet all hosts experience significant exponential growth and decline of plasma viral load during acute infection [1]. We chose a representative value of *b*_tot_ = 6 day^−1^ for our population dynamics simulations.

#### The mutation rate, *μ*

The overall rate at which nucleotide substitutions occur during HIV infection was estimated at 1. 2 × 10^−5^/nucleotide/day [16], and more specifically varies between each pair of nucleotides [6, 16]. Here, we consider a simplified probability of mutation *μ* for binary amino acid residues that is three times the nucleotide substitution rate, *μ* = 3.6 × 10^−5^/site/generation. However, in S1 Appendix we extend our methods to allow for site and state-dependent mutation rates, compute the transition probabilities *μ*_*i*,0→1_ and *μ*_*i*,1→0_ for the patient used in the example below, and show that the resulting dynamics of mutations are similar to the main text. Constructing a full amino acid substitution matrix using the codon map requires a model with multiple states per site, which is beyond the scope of this paper.

Table 1 summarizes all of the parameter values we used.

**Table 1.** **Parameter values used for the application of EMF to the dynamics of p24.**

| parameter | description | value (units) |
| --- | --- | --- |
| $L$ | length of consensus p24 amino acid sequence | 231 sites |
| $\beta$ | proportionality constant between prevalence and fitness landscapes: $h_i = \beta h_i^p$, $J_{ij} = \beta J_{ij}^p$ | $0.023 \text{ day}^{-1}$ |
| $F_0$ | intrinsic fitness of consensus sequence | $1.6 \text{ day}^{-1}$ |
| $b_{\text{tot}}$ | overall magnitude of CTL response | $6 \text{ day}^{-1}$ |
| $t_0$ | time delay before activation of CTL response | 7 days |
| $t_{1/2}$ | time to half-maximum of CTL response | 7 days |
| $n_{\text{Hill}}$ | Hill coefficient of CTL response | 2 |
| $\mu$ | mutation rate | $3.6 \times 10^{-5} \text{ site}^{-1} \text{ day}^{-1}$ |
| $N_0$ | initial population size | 10 |

<sup>2</sup>We found that HIV dynamics do not depend sensitively on the precise form of $b_\varepsilon(t)$ , and in the Discussion we propose an extension of our methods that include explicit consideration of CTL clone populations.

#### Patient CH58 [3]

Here we describe a patient (data taken from [3]) whom we use as an example for application of the EMF method. Patient CH58 was a male infected by an HIV-1 group M subtype B strain [3]. He was not treated with antiretroviral therapy (ART) during the period of study, in accordance with contemporaneous medical protocol. Blood plasma was drawn at several timepoints from which snapshots of the intra-host HIV population were determined by single genome amplification and sequencing [8], and the specificity and magnitude of HIV-specific CTL responses were also mapped by interferon-γ ELISpot assays against overlapping peptides spanning the founder viral sequence [3]. The founder sequence had five mutations in p24 w.r.t. the subtype B consensus, and two p24 epitopes were targeted by the patient (see Fig 5(A), left panel; one epitope in Env and one in Nef were also targeted (not shown)). Mutations away from the founder sequence in each blood sample are listed in Fig 5(A), right panel. Frequencies of mutations in the two p24 epitopes over time are also plotted in Fig 5(B).

## Results

### Application of EMF to predict p24 mutational dynamics in patient CH58

As a proof of principle, we apply the EMF method to simulate the dynamics of the p24 gag protein within a patient whose CTL responses and dynamics are known (taken from [3]). Note that the main goal of this section is to demonstrate use of the method, and not to prove that simulated HIV dynamics using fitness landscapes can predict with good accuracy the locations and timescales of escape mutations (as was done in [24] for a larger number of HIV proteins and patients).

As input to EMF, we use:

1. the binarized p24 founder sequence infecting patient CH58 (Fig 5(A), left panel);
2. the within-host fitness landscape of p24 (Eq 2), with parameters *F*_0_, {*h*_*i*_} and {*J*_*ij*_} inferred in Parameter estimation, the locations of the p24 epitopes (Fig 5(A), left panel), and their magnitudes *b*_1,0_ and *b*_2,0_, where *b*_1,0_ + *b*_2,0_ = *b*_tot_.

In the following, we first consider *b*_1,0_ = *b*_2,0_, and study the effect of the vertical immunodominance, *b*_1,0_ ≠ *b*_2,0_, later.

### EMF outputs coupled dynamics of effective fields and frequencies of mutation at each HIV residue over time

Starting with the 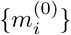 in Fig 5(A), left panel, we follow Eq (19) to recursively compute the *L* effective fields 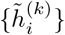 and frequencies of mutations 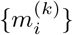 over time *k*, which are plotted in Fig 6. We find that following the activation of the CTL response (yellow dotted lines), the effective fields at sites within epitopes rise above zero (Fig 6(A)), signifying a selective pressure to mutate to *s*_*i*_ = 1 at these sites. At longer times, two sites (15 and 110 as shown later) have large 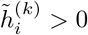, while the rest are distributed around a value less than zero.

The effective fields dictate the dynamics of the mutational frequencies 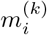 (Fig 6(B)). Positive 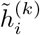 lead to increasing 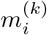, while negative 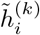 lead to decreasing 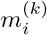, with the magnitude of 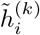 determining the rate of change. In turn, changes in the 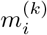 feed back into the time-varying effective fields 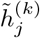 (for *j* ≠ *i*) through terms depending on *J*_*ij*_ and *b*_*ε*_ in Eqs (14) and (16), thus prescribing how the changing sequence background modifies the tendency for each HIV residue to mutate.

**Fig 5.**
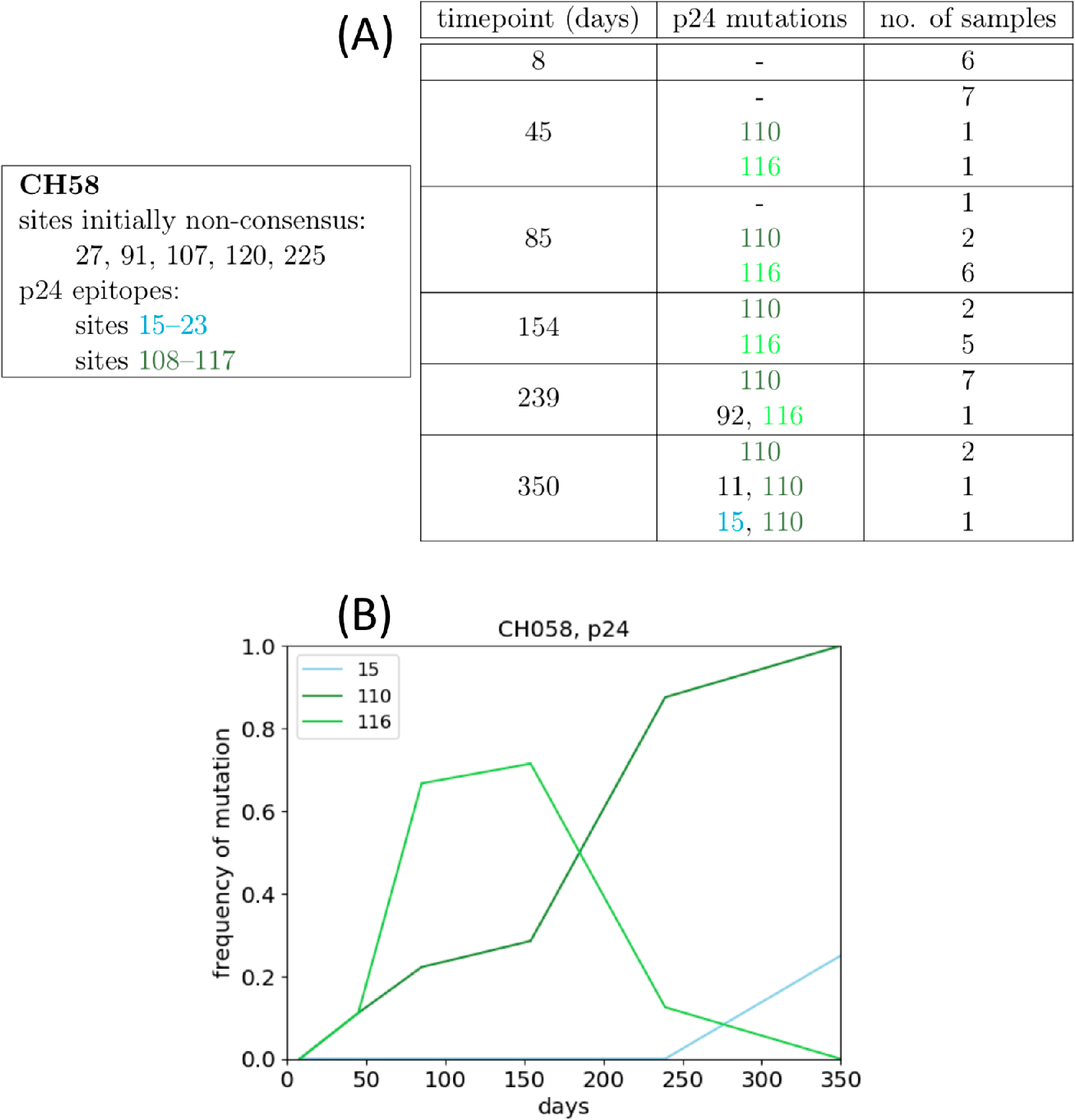
Dynamics of p24 measured in patient CH58 (data taken from [3]) (A) Left panel: Non-consensus residues in the founder sequence and epitopes in p24 targeted by patient CH58. Right panel: p24 mutants and sampled frequencies drawn from patient CH58 at multiple timepoints post-infection. Each row represents a distinct sampled sequence. (B) Frequencies of mutations at sites 15, 110 and 116 sampled from patient CH58 over time.

### EMF combines fitness costs and epistatic interactions to predict the locations of HIV escape mutations

Focusing on the two p24 epitopes, we find that mutations initially arise at all sites within each epitope, but eventually one fixes while the others decline (Fig 7). This behavior may be discerned from the EMF equations: if the frequency of mutation 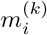 at site *i* in an epitope is close to 1, the last term of Eq (16) will be small for sites *j* ≠ *i* in the epitope, but not for site *i*. There is thus a tendency for the other 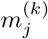 to decline to zero, leaving all of the weight of *b*_*ε*_(*k*) to act on site *i* and keeping it mutated.

EMF combines fitness costs and epistatic interactions to determine which escape mutations arise. In the first p24 epitope, site 15 is the least-fitness-cost mutation according to the intrinsic landscape *F*_in_(*S*) (Fig 7(A) inset) and is the one that fixed. In the second epitope, site 116 is the least-fitness-cost mutation (Fig 7(B) inset) but site 110 fixed, and indeed 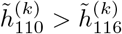 at all times *k* (not shown). In the patient, an escape mutation at site 110 fixed by 350 days post-infection (Fig 5(B)), and furthermore there appeared to be competition between mutations at sites 110 and 116 at intermediate times (Fig 5(B)), qualitatively resembling the dynamics produced by EMF for this patient (Fig 7(B)). We emphasize that this is merely one example that we chose to present, but Barton et al. [24] showed empirically (by performing 10^3^ stochastic simulations per HIV protein and host) that HIV fitness landscapes inferred from global prevalence provide good predictions of the locations of HIV escape mutations, for a larger number of patients and proteins studied [24]. Here, we have shown that EMF is capable of making similar predictions that are directly encoded in the effective fitnesses.

**Fig 6.**
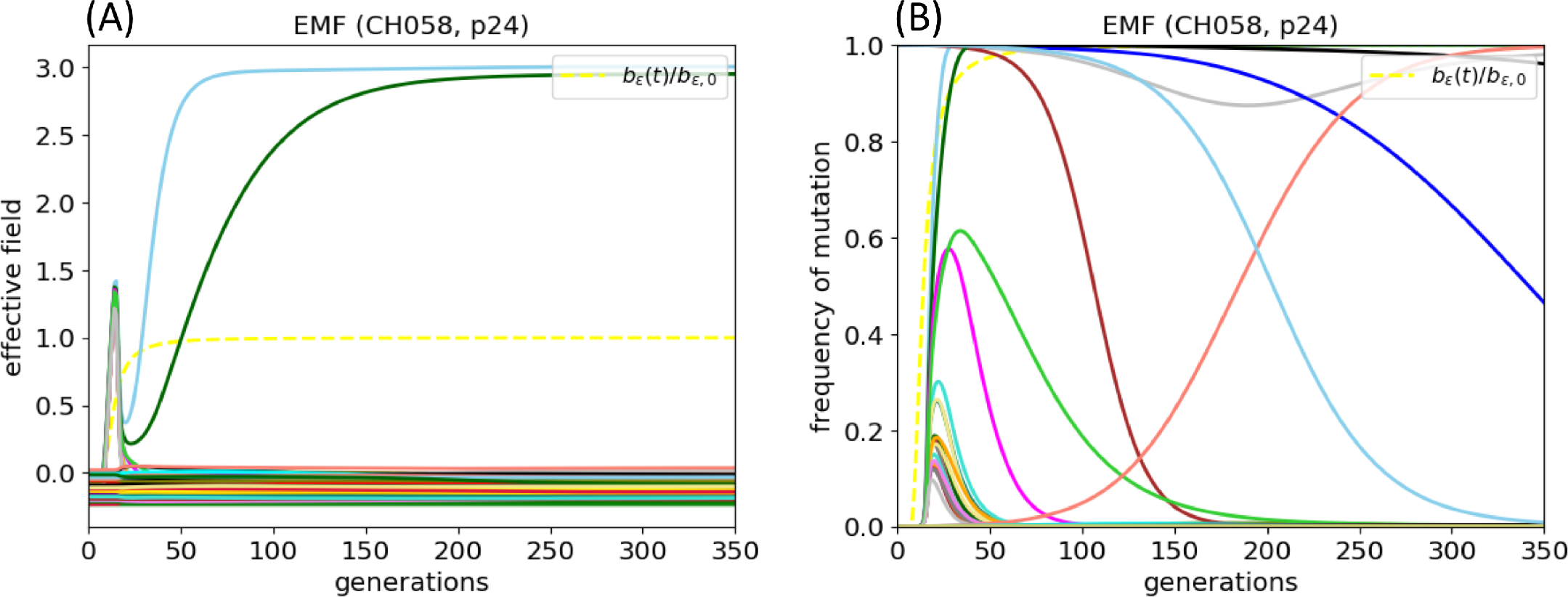
Dynamics of effective fields and frequencies of mutations output by EMF. (A) Effective fields 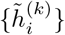 and frequencies of mutations 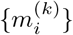, at each of the *L* residues of p24 during evolution within patient CH58, as predicted by the EMF method. Plotted in the yellow dotted lines are the CTL response *b*_*ε*_(*t*)/*b*_*ε*,0_ (Eq (22)). One generation of EMF corresponds to 1 day because we have used parameter values in units of day^−1^ (see Table 1).

**Fig 7.**
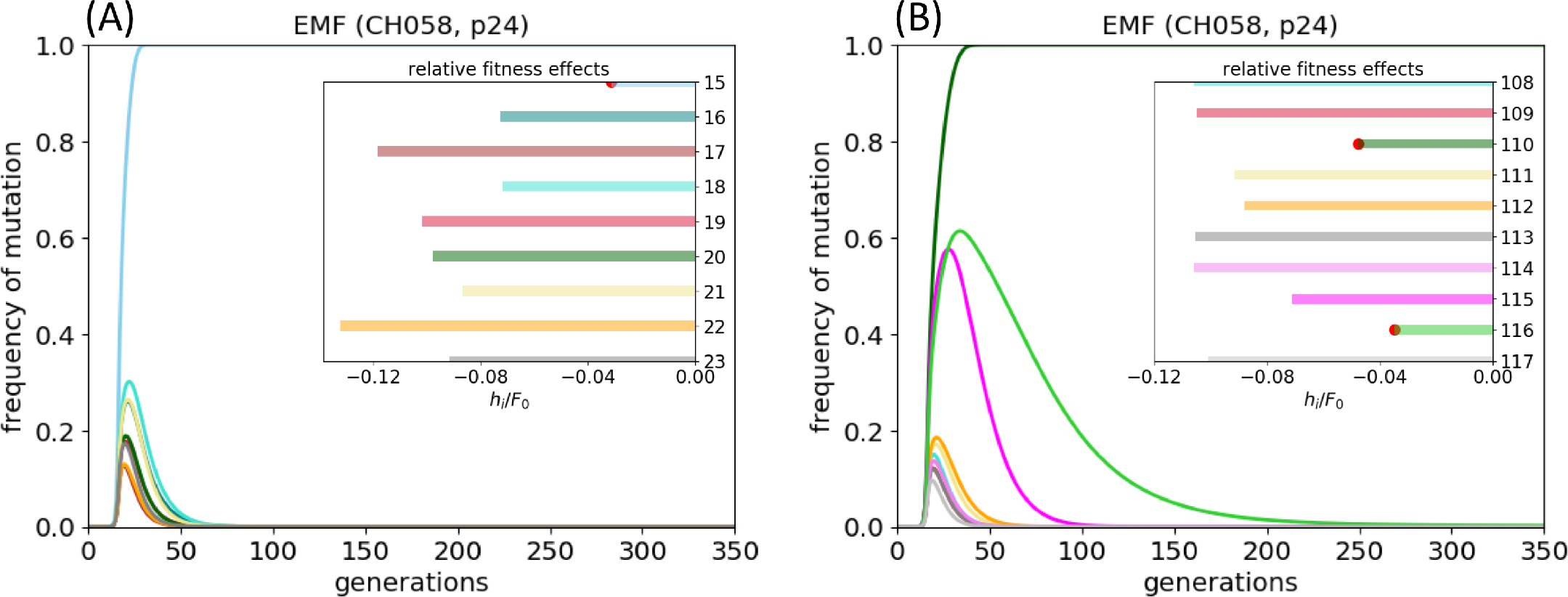
Frequencies of mutations in p24 epitopes targeted by patient CH58. Frequencies of mutations in the p24 epitopes at (A) sites 15–23, and (B) sites 108–117, as predicted by the EMF method. Insets show *h*_*i*_/*F*_0_ at each site within each epitope, and the least-fitness-cost mutations are marked with red circles.

### EMF predicts the vertical immunodominance of CTL responses to affect the order and timescale of escape mutations by modifying the relative strengths of effective fitnesses

In patient CH58, a mutation at site 15 was only detected on day 350 (Fig 5(B)). This delay might be caused by a smaller CTL population targeting the first p24 epitope, and hence a weaker selective pressure for sites 15–23 to mutate. Indeed, the peak CTL response at this epitope was measured by ELISpot to be smaller by a factor of 4–5 than at sites 108–117 [3]. To study how EMF predicts the vertical immunodominance of CTL responses to change the dynamics of mutations, we repeated the above simulation using *b*_1,0_ = *b*_2,0_/5 and keeping the same overall *b*_tot_ = *b*_1,0_ + *b*_2,0_ as before. The resulting dynamics are shown in Fig 8(A) (solid lines). The same escape mutations arise, but unlike the *b*_1,0_ = *b*_2,0_ case (dotted lines), escape at site 15 occurs later than at site 110. This delay would be compounded by a larger time lag *t*_0_ for the first epitope as compared with the second, as was observed in the patient [3] (not shown). Indeed, Liu et al. [3] performed a statistical analysis for a larger number of patients and proteins and showed that the vertical immunodominance correlates well with the rate of HIV escape. Here, EMF encodes this effect simply by modifying the relative strengths of the effective fitnesses.

**Fig 8.**
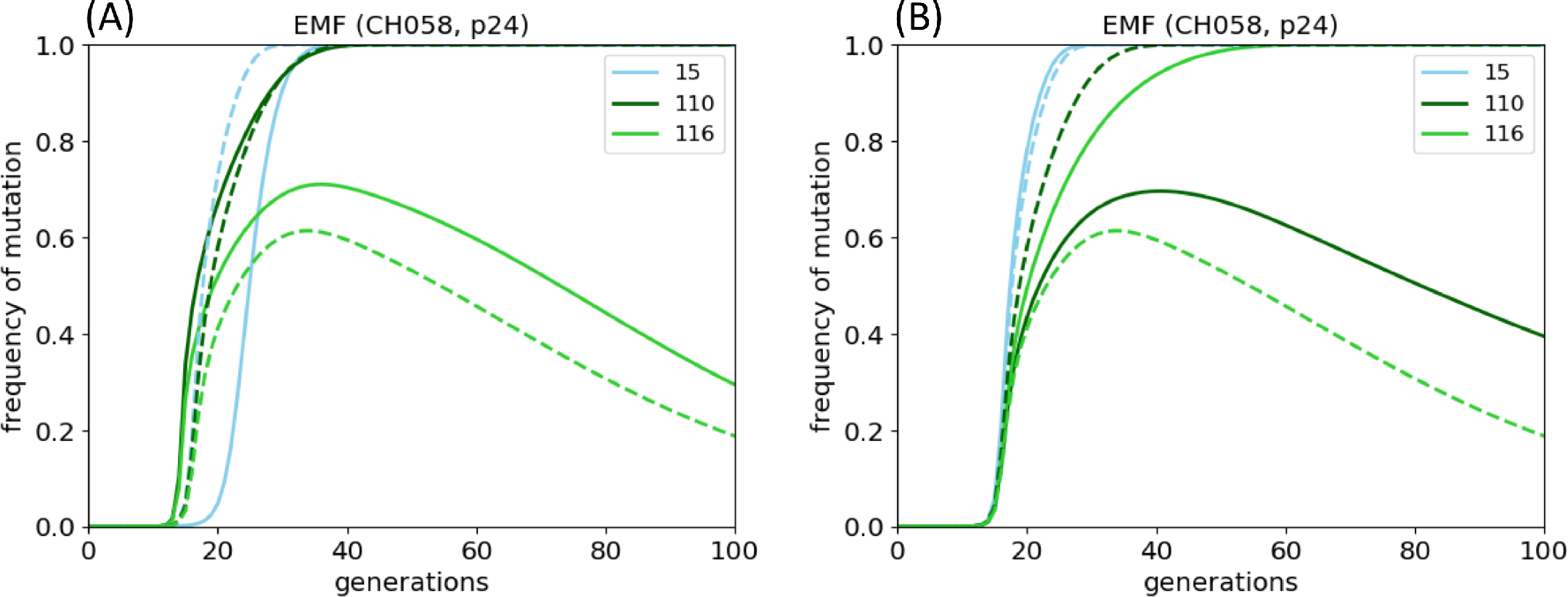
Effect of vertical immunodominance and sequence background on the dynamics of p24 escape mutations in patient CH58. (A) Mutational dynamics at sites 15, 110 and 116 when *b*_1,0_ = *b*_2,0_/5 (solid lines), and *b*_1,0_ = *b*_2,0_ (dotted lines), for the same overall *b*_tot_ = *b*_1,0_ + *b*_2,0_. (B) Mutational dynamics at sites 15, 110 and 116 when the founder sequence is the NL4-3 strain (*s*_120_ and *s*_208_ = 1; solid lines), as compared with the actual founder sequence of patient CH58 (dotted lines). Here, site 116 mutates instead of 110.

### EMF predicts that a different sequence background may cause different escape mutations to arise in a host because of compensatory interactions

In models of HIV dynamics with multiple HIV strains that do not consider epistatic interactions (e.g. [25–28, 32]), the sequence background outside of an epitope does not affect the locations of escape mutations in the epitope. Here, we asked if a different founder sequence might affect the locations of escape mutations, for a host that mounts the same CTL responses as patient CH58. We simulated the dynamics starting with, as an example, the NL4-3 strain (used in the experiments of [13], with mutations at sites 120 and 208 in p24 w.r.t. the subtype B consensus). Fig 8(B) (solid lines) shows the result for sites 15, 110 and 116. In contrast with the above (dotted lines), site 116 acquired an escape mutation instead of 110, which is in fact the naïve expectation based on knowledge of just the fitness costs (Fig 7(B) inset). Thus, knowing the epistatic interactions of the fitness landscape may enable more accurate predictions of which escape mutations arise given the sequence background, which were indeed the findings of [24] for a larger number of HIV proteins and patients studied. Here, we have shown that EMF efficiently makes such predictions that are encoded in the effective fitnesses.

### EMF predicts reversions and compensatory mutations occurring over longer timescales

Apart from mutational escape within epitopes, EMF also informs the dynamics of mutations outside of epitopes. Figure 9 shows the effective fields and frequencies of mutations at all p24 residues outside of the two epitopes output by EMF for patient CH58. The effective fields are distributed around a value less than zero and are mostly negative. Because their magnitudes are closer to zero than at sites within epitopes (Fig 6(A)), the mutational dynamics they induce tend to occur over longer timescales. We find that the initially non-consensus sites revert back to consensus over a range of timescales (Fig 9(B)), which is qualitatively consistent with the findings of Refs [10, 11] and especially [9] that found that sites tend to revert to the group or subtype consensus residue throughout the course of intra-patient evolution in all of the patients studied. Within the EMF framework, a distribution of reversion timescales is due to a distribution of effective fitnesses that are negative and close to zero.

**Fig 9.**
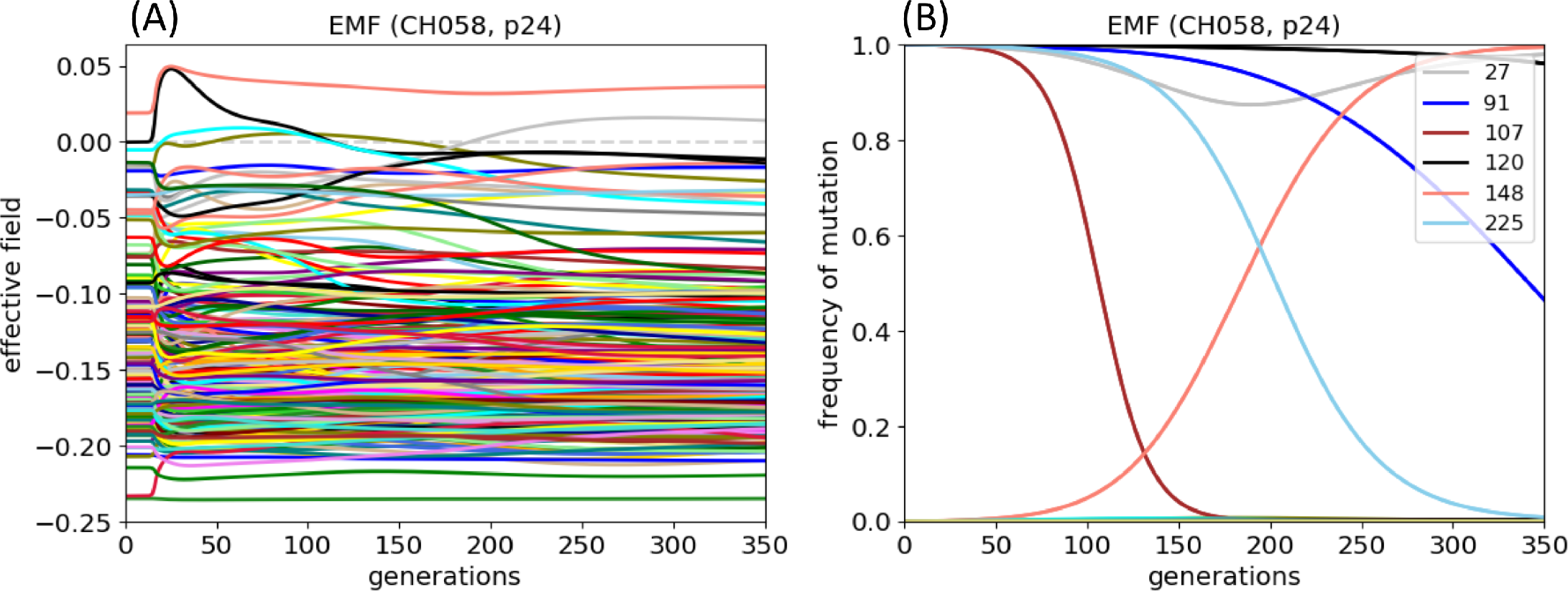
Reversions and compensatory mutations in patient CH58, as predicted by EMF. (A) Effective fields 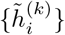, and (B) frequencies of mutations 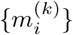, at p24 residues outside of epitopes for patient CH58, as predicted by the EMF method. Compensatory mutations occur at sites whose effective fields rise above zero (here, site 148). Reversions occur at sites initially non-consensus whose effective fields remain negative (here, sites 91, 107, 120, and 225). Site 27 was initially non-consensus but not predicted by EMF to revert.

Finally, EMF predicts 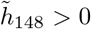 at long times (Fig 9(A)) and hence a potential compensatory mutation to arise at site 148 (Fig 9(B)). This did not occur in the patient; instead, mutations at sites 92 and 11 arose (Fig 5(A), right panel). We emphasize that there are many possible reasons why mutations predicted by EMF may not occur in the patient, and vice-versa: EMF is an approximate high-recombination-rate model that ignores many important processes during HIV infection, a real infection presumably has large stochastic effects that cannot be precisely predicted, the fitness landscape may be incomplete, etc. (see Discussion). For patient CH58, the site 92 mutation appeared with the site 116 mutation on day 239 at a low frequency before the site 116 mutation went extinct, so the site 92 mutation could possibly be a hitchhiker mutation, which cannot be modeled by EMF. The site 11 mutation might also affect antigen processing of the epitope at site 15–23 and hence its presentation to CTLs [46, 47], allowing evasion of CTL pressure despite not having a mutation within the actual epitope; indeed it arose at the same time as the site 15 escape mutation. Our current method does not account for antigen-processing mutants, although *F*_host_ can easily be modified to do so.

### Mean fitness and site entropy of the intra-host population

The mean fitness and other quantities describing the entire intra-host population are particularly simple to compute within the EMF framework, which we show here. The mean fitness at time *k* is given by

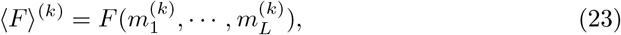

which is plotted in Fig 10(A) for *F* = *F*_host_ (blue curve) and *F* = *F*_in_ (orange curve). As host CTLs are activated, the founder sequence becomes strongly selected against, which is represented by the valleys in Fig 10(A). The intra-host population evolves over time to “climb” out of this fitness valley through the generation of mutants and selection of beneficial ones. Thus, the mean fitness first decreases, and then is a strictly nondecreasing function of time. The blue and orange curves merge because for sequences that have completely escaped host CTL responses, *F*_host_ and *F*_in_ are identical.

**Fig 10.**
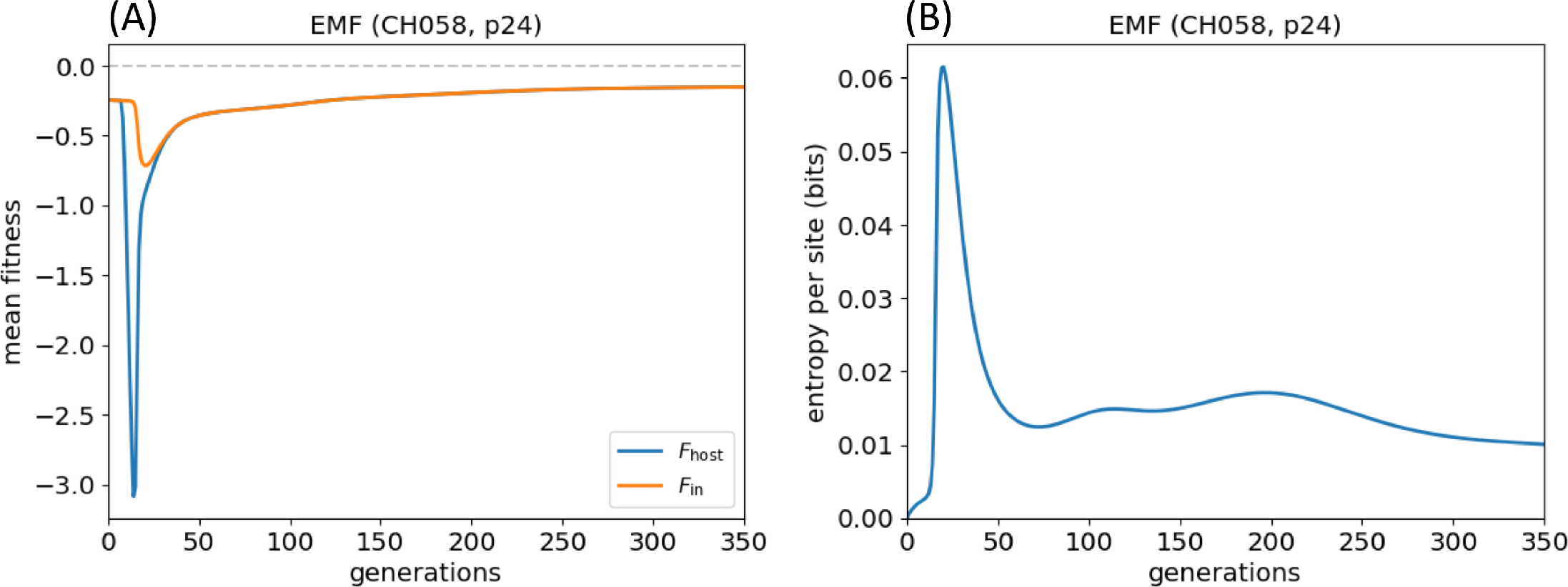
Mean fitness and site entropy of the intra-host population in patient CH58 over time, as predicted by EMF. (A) Mean fitness 〈*F*〉^(*k*)^ − *F*_0_ of the intra-host population over time, using *F* = *F*_host_ (blue line) and *F* = *F*_in_ (orange line). The grey dotted line is the fitness of the consensus sequence (0, …, 0). (B) The mean entropy per site over time.

A measure of genetic diversity of the intra-host population is the mean entropy per site,

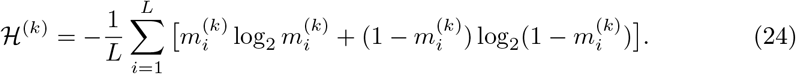

This quantifies how different the frequencies of mutation are, on average, from 0 or 1 (−*m*log_2_*m* − (1 − *m*)log_2_(1 − *m*) is bounded by 0 and 1, equals 0 at *m* = 0 and 1, and equals 1 at *m* = 1/2). Figure 10(B) shows that the mean site entropy spikes when several escape mutations arise simultaneously when CTL responses are activated, and declines as escape mutations either fix or disappear. Even at long times, the mean site entropy remains at a finite value because mutations are continuously generated at all sites but remain at low levels due to being less fit; this is a manifestation of mutation-selection balance [38]. (Consistently, Zanini et al. [16] used mutation-selection balance to infer the fitness costs at all sites in the HIV genome not undergoing CTL-driven selection.)

### Application of EMF stochastic population dynamics method

In Methods, we introduced a stochastic population dynamics method based on the EMF approach, where population size changes depend on the frequencies of mutations 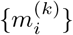 in the population at each time, unlike prior stochastic simulation approaches with a fixed population size [24, 32] or a time-varying one that is externally defined [33]. In the following, we apply this method again to the dynamics of p24 gag in patient CH58, assuming the initial population size to be *N*_0_ = 10, and keeping all other parameter values the same as before (see Table 1).

### EMF population dynamics simulations produce variability in the locations of escape mutations, but are consistent with the deterministic EMF method

Figures 11(A)–11(C) show the dynamics of mutations at sites 15, 110 and 116 in ten representative stochastic runs (dotted lines), their mean over 200 stochastic runs (dashed lines), and a comparison with the deterministic EMF method above (solid lines). Figures 11(D)–11(E) show the mean of 200 stochastic runs (dashed lines) and the deterministic dynamics (solid lines) at all sites in the two p24 epitopes. Figures 11(F)–11(G) show in greater detail the fraction among 200 stochastic runs where a site in each of the epitopes has the highest frequency of mutation at the end of the simulation.

We find that the stochastic method produces escape events in individual runs that can be very different from the deterministic EMF prediction: with some probability, other sites may fix, particularly when they happen to arise early enough in a simulation (not shown). This is why the mean values for sites 15, 110 and 116 are lower than in the deterministic case, and the averages for other within-epitope sites are higher (see Figs 11(D)–11(E)). However, for both epitopes, the deterministic EMF method is a good predictor of the stochastic case averaged over many simulation runs. In the first epitope, site 15 fixes most of the time (Fig 11(F)), and in the second, greater variability in the locations of escape mutations (particularly at sites 115 and 116; see Fig 11(G)) seem to correspond with higher frequencies of mutation at these sites at intermediate times in the deterministic dynamics (Fig 11(E)). Hence, these simulations suggest that adding stochasticity to our methods produces variability in the dynamics of escape, but on average, the results are consistent with the deterministic case.

**Fig 11.**
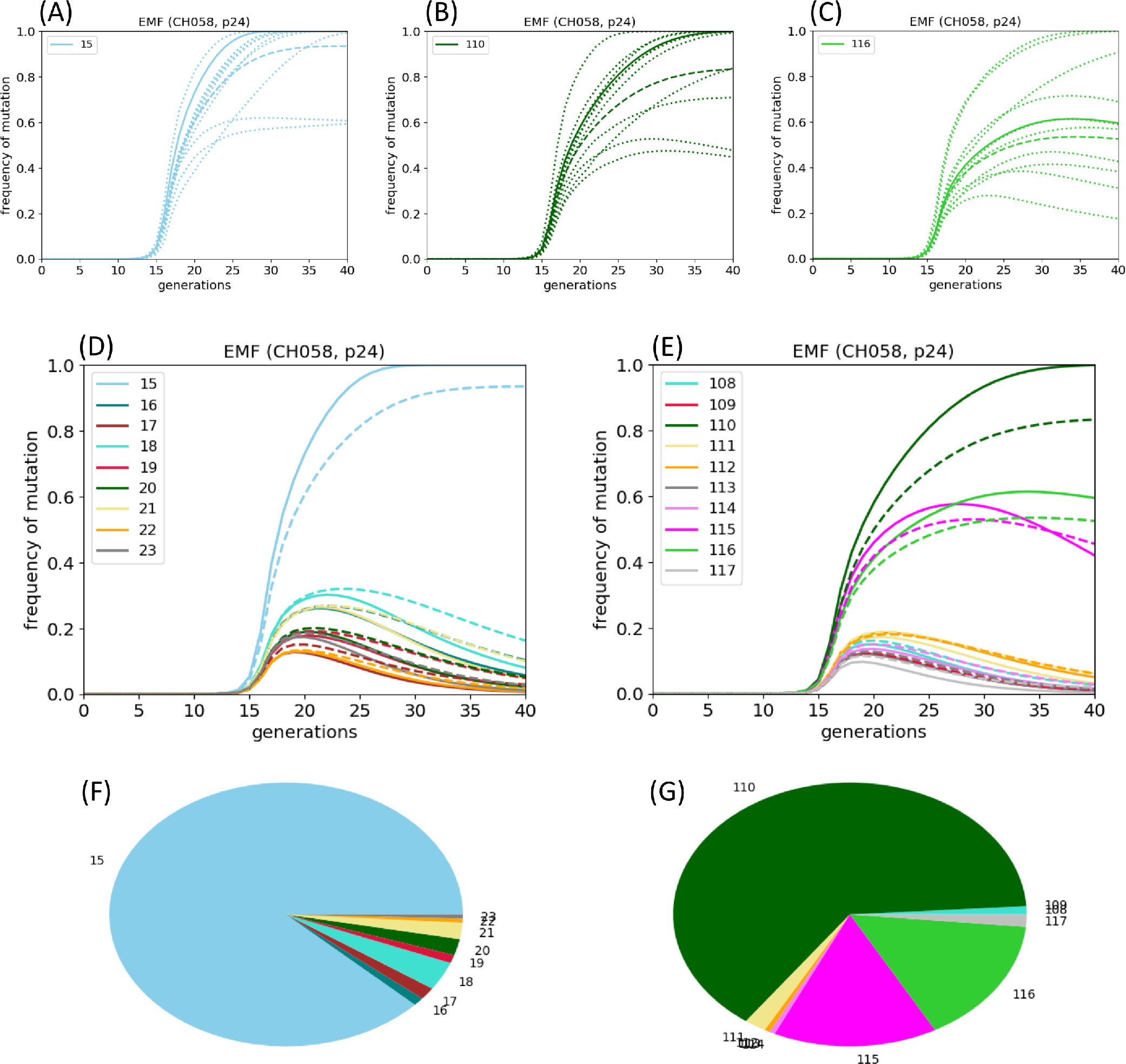
Stochastic dynamics of escape as predicted by EMF population dynamics simulations. Frequencies of mutation at (A) site 15, (B) site 110, and (C) site 116 produced by the deterministic EMF method (solid lines), 10 representative stochastic runs (dotted lines), and the mean over 200 independent stochastic runs (dashed lines). (D)–(E) Frequencies of mutations within the two p24 epitopes produced by the deterministic EMF method (solid lines), and the mean over 200 independent stochastic runs (dashed lines). We did not plot error bars because the dynamics lead to sites having mutational frequencies tending to 0 or 1, so the spread around the mean does not reflect the results of individual stochastic runs. (F)–(G) Fraction among 200 stochastic runs where a site has the highest frequency of mutation in the epitope at the end of the simulation. Site 15 fixes most of the time in the first epitope, but there is greater variability in the second, especially at sites 115 and 116 instead of 110, which is consistent with the deterministic EMF dynamics.

### EMF population dynamics simulations produce exponential growth and decline of the population size, consistent with viral load kinetics during acute infection

Figure 12 shows the log population size over time for the determinstic case (blue line), and 10 stochastic simulation runs (red lines). The population first increases exponentially, then exponentially declines following activation of CTL responses. These dynamics are consistent with plasma viral load kinetics in untreated hosts [1], and also experiments that found a rapid rebound in plasma viral load following CTL depletion that reversed upon the replenishment of these cells [5]. Furthermore, this characteristic exponential growth and decline is observed for a range of *b*_tot_ (see S1 Fig), which is consistent with the fact that diverse hosts, who presumably have a range of magnitudes and specificities of CTL responses, experience qualitatively similar viral load kinetics during acute infection.

**Fig 12.**
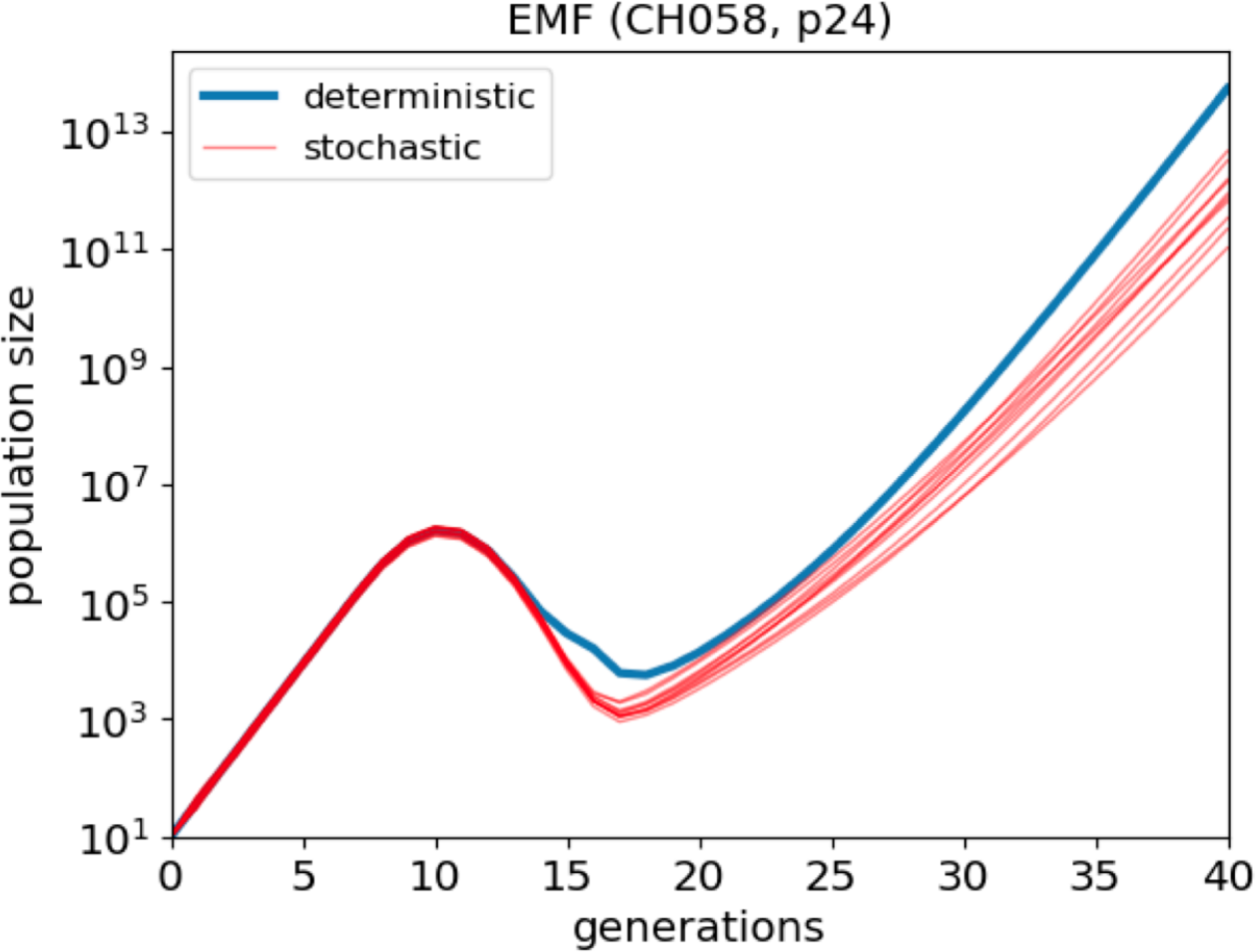
Dynamics of the population size in patient CH58 using EMF population dynamics simulations. Dynamics of population size over time for the deterministic EMF method (blue line) and 10 representative stochastic runs (red lines). The exponential increase and initial decrease in the population size qualitatively resemble plasma viral load kinetics during early acute infection. However, the unbounded exponential growth after viral escape clearly does not reflect a real infection, and is a result of not explicitly considering viral infection and elimination of target cells.

While the exponential increase in viral load and initial decline after activation of host CTL responses resemble viral load kinetics in a real infection, the intra-host population quickly overcomes host CTL pressure and continues to grow exponentially. In actual patients, this does not happen, for two important reasons:

1. We did not consider an explicit target cell population that declines following infection. Models that include the dynamics of host target cells via ordinary differential equations show an approach to a finite fixed point as opposed to unbounded growth (see e.g. [28]). In the Discussion, we propose how our methods may be extended to include dynamics of an explicit target cell population.
2. There are immunodominance shifts in the CTL response. CTLs specific for epitopes from which the HIV population has escaped decline, and new CTL clones specific for other epitopes emerge (see e.g. [25]). Thus, there is continuous CTL selection and HIV escape throughout the course of intra-host infection, which we did not attempt to model here.

## Discussion

In this paper, we introduced an approach for simulating HIV dynamics given a fitness landscape, which we designated the evolutionary mean-field (EMF) method. EMF is an approximate high-recombination-rate model of HIV replication and mutation, with time-varying population sizes. EMF takes as input the fitness landscape of an HIV protein, including epistatic interactions, and the locations and strengths of a host’s CTL responses, and outputs time-dependent “effective fitnesses” describing the tendency for each HIV residue to mutate, and frequencies of mutations caused by these time-varying fitness effects. Applying this method to the dynamics of the p24 gag protein in a patient whose CTL responses are known (from [3]), we showed how fitness costs and compensatory interactions, vertical immunodominance of CTL responses, and the sequence background modify the effective fitnesses and hence impact the locations and relative timescales of HIV escape mutations, which is consistent with previous work employing stochastic simulations [24]. These include cases where knowledge of the epistatic interactions improves upon predictions relying simply on fitness costs. We also show that features of longer-term dynamics, specifically reversions, may be described in terms of the effective fitnesses, which is also qualitatively consistent with other work [9]. EMF makes quick predictions that may otherwise require performing many stochastic simulation runs [24], and furthermore describes various genetic-level attributes known to influence HIV dynamics in terms of their combined effect on the effective fitnesses.

We also developed a stochastic population dynamics method based on EMF, and quantified the variability in the escape mutations that arise in the same example. The HIV population sizes resulting from this method show a characteristic exponential rise and fall during early infection, consistent with observed viral load kinetics, although the unbounded exponential growth following HIV escape is a weakness of the current implementation, and we suggest below extensions to overcome it.

What have we lost from a high-recombination-rate approximation of HIV dynamics?

Unlike a real HIV infection, here escape mutations in separate epitopes quickly recombine, forming variants that escape multiple epitopes; in this sense EMF simulates a “stronger” virus. Also, EMF is unable to account for important features such as clonal interference and genetic hitchhiking that do not occur in linkage equilibrium. On the other hand, EMF provides quick dynamical predictions that can be checked against simulations implementing a finite recombination rate [24], and we expect that, averaged over many hypothetical runs, mutations arising due to fitness effects in a model with finite recombination rate should be comparable and consistent with the results of EMF. Indeed, in S2 Appendix we performed this check and validated the results obtained in the above example with Wright–Fisher simulations with a finite recombination rate (as in [24]). Neher and Shraiman [31] obtained analogous mean-field equations to Eq (14) as the lowest order in an expansion in the inverse recombination rate; it would be interesting to consider the next-order corrections (i.e., quasi-linkage equilibrium) which would be a finite-recombination-rate extension of EMF (this would however require *L*(*L* −1) extra equations, i.e. 5 × 10^4^ for p24).

We have presented EMF as a discretized-time algorithm where each time step represents one day (see Table 1), which roughly corresponds to a replication cycle of HIV [34]. We could of course discretize time more finely, scaling the replication and mutation rates appropriately, i.e., if one time step represents *κ* ≪ 1 days, then *F*_0_ → *κF*_0_, *b*_tot_ → *κb*_tot_, and *β* → *κβ*. In this limit, the HIV population size obeys

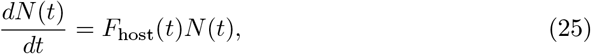

where *F*_host_(*t*) depends on the genetic composition of the population at time *t*, and CTL pressure is implicit in *F*_host_. Thus, mutational escape leads to continued exponential growth (Fig 12). To overcome this limitation, we may consider target cell and CTL clone populations explicitly (much like in other compartmental models of HIV dynamics [25–28]):

1. To account for explicit viral infection of target cells and target cell dynamics, we propose extending Eq (25) to a pair of coupled equations:

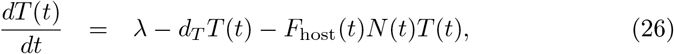

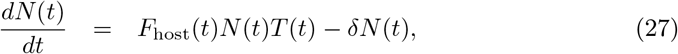

where *λ* and *d*_*T*_ are the influx and natural death rates of target cells, and *δ* is the rate of clearance of viral particles. Effectively, these equations extend two-compartment models of HIV dynamics (see e.g. [28]) to have viral infectivity that depends on the genetic composition of the intra-host population. Models that include target cell dynamics produce viral population sizes that approach a finite fixed point [28]; in essence, “fit” viruses do not grow unboundedly if target cells are limiting. Within the EMF procedure, we propose first computing an intermediate 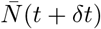 using Eq (20), which accounts for population size changes due to the fitness of strains in the population, and then *T*(*t* + *δt*) and *N*(*t* + *δt*) using Eqs (26) and (27) (with 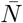 in place of *N* on the RHS of these equations).
2. To account for explicit CTL clone dynamics, we propose extending Eq (25) to have one equation for each CTL clone:

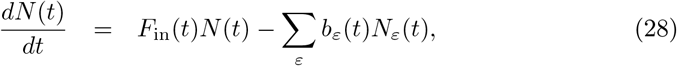

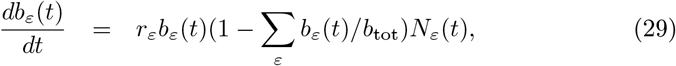

where Eq (28) uses *F*_in_ instead of *F*_host_, *r*_*ε*_ is the growth rate of the CTL population specific for epitope *ε*, *N*_*ε*_(*t*) is the HIV subpopulation susceptible to these CTLs, and *b*_tot_ is now a carrying capacity. While this paper developed an approach to HIV dynamics *given* a fitness landscape, the method is presumably limited by the quantitative accuracy of the landscapes used. While we have limited ourselves to binary sequences, using fitness landscapes of nucleotide or amino acid sequences (such as those inferred in [13, 24]) would likely improve the accuracy of EMF’s predictions. Considering higher-order interactions in the landscape would also help, should these become known. Extensions of EMF to non-binary sequences and to fitness landscapes with higher-order interactions are straightforward, but beyond the scope of this paper. Also, we estimated the p24 fitness landscape from its prevalence in the global population of hosts (following [12, 13, 24]), but quantitative estimates of fitness from prevalence are limited by several issues as studied previously [14–16, 37]; in particular, prevalent HIV mutations are more likely to be HLA-associated than less prevalent ones [16]. Alternatively, measurements of intrinsic fitness from cell culture experiments [14, 15], including epistatic interactions, may eventually lead to more accurately known fitness landscapes of HIV proteins. This may lead to more accurate predictions of the intra-host dynamics computed using EMF, and other methods.

## Supporting information

**S1 Appendix. EMF with state and site-dependent mutation rates.**

**S2 Appendix. Validation of EMF dynamics results by Wright–Fisher simulations.**

**S1 Fig. EMF population dynamics produces a characteristic exponential rise and fall of population size during the early stages of infection, for a range of** *b*_tot_.

## Acknowledgments

We thank Arup K. Chakraborty and John P. Barton for insightful discussions and a close reading of earlier versions of the manuscript. We also thank John P. Barton for help with inferring the p24 prevalence landscape. HC was supported by an A*STAR Scholarship and the Ragon Institute of MGH, MIT and Harvard. MK acknowledges support from NSF through grant number DMR-1708280.

## Footnotes

1 Note that our methods can easily be extended to fitness landscapes with higher-order interactions; indeed, we model CTL epitopes as a higher-order interaction (see later).

2 We found that HIV dynamics do not depend sensitively on the precise form of *b*_*ε*_(*t*), and in the Discussion we propose an extension of our methods that include explicit consideration of CTL clone populations.

