## Supplementary material for "Mean-field computational approach to HIV dynamics on a fitness landscape": S1 Appendix

### EMF with state and site-dependent mutation rates

The binary approximation of HIV sequences involves replacing the consensus amino acid at position $i$ with 0 and any other state (amino acid or gap) at position $i$ with 1. Because the substitution rates between different pairs of nucleotides are different [1,2], and because there is a mapping between codons and amino acids, within the binary approximation the amino acid mutation rates should in general be different between $0 \to 1$ and $1 \to 0$ and for different sites $i$. In this Appendix, we describe how the EMF equations are modified as a result, and apply them again to the dynamics of p24 infecting patient CH58.

#### Modifications to EMF

First, let us consider a one-site model where $s_i = 0$ has fitness $F_0$ and $s_i = 1$ has fitness $F_0 + h_i$. Also suppose that $s_i = 0$ mutates to $s_i = 1$ with probability $\mu_{i,0\to1}$/generation and $s_i = 1$ mutates to $s_i = 0$ with probability $\mu_{i,1\to0}$/generation. Suppose the frequency of $s_i = 1$ in a population of size $N_i^{(k)}$ at time $k$ is $m_i^{(k)}$. The population sizes of $s_i = 0$ and $s_i = 1$ at time $(k+1)$ are thus

$$\begin{pmatrix} e^{F_0} & 0 \\ 0 & e^{F_0+h_i} \end{pmatrix} \begin{pmatrix} 1-\mu_{i,0\to1} & \mu_{i,1\to0} \\ \mu_{i,0\to1} & 1-\mu_{i,1\to0} \end{pmatrix} \begin{pmatrix} 1-m_i^{(k)} \\ m_i^{(k)} \end{pmatrix} N_i^{(k)}$$
$$= e^{F_0} \begin{pmatrix} (1-\mu_{i,0\to1})(1-m_i^{(k)}) + \mu_{i,1\to0} m_i^{(k)} \\ e^{h_i}[\mu_{i,0\to1}(1-m_i^{(k)}) + (1-\mu_{i,1\to0}) m_i^{(k)}] \end{pmatrix} N_i^{(k)}, \tag{1}$$

giving a fractional population size change at time $(k+1)$ of

$$\tilde{Z}_i^{(k+1)}(m_i^{(k)}) \equiv \frac{N_i^{(k+1)}}{N_i^{(k)}} = e^{F_0}\{(1-\mu_{i,0\to1})(1-m_i^{(k)}) + \mu_{i,1\to0} m_i^{(k)}$$
$$+ e^{h_i}[\mu_{i,0\to1}(1-m_i^{(k)}) + (1-\mu_{i,1\to0}) m_i^{(k)}]\}. \tag{2}$$

For sequences of length $L$ replicating according to a fitness landscape $F = F_{\text{host}}$, the number of offspring with sequence $S^{(k+1)}$ produced by a single sequence $S^{(k)}$ in one generation is given by

$$\langle S^{(k+1)}|T|S^{(k)}\rangle = e^{F(S^{(k+1)})} \prod_{i=1}^{L} \left[ (1-\mu_{i,0\to1})^{\delta_{s_i^{(k)},s_i^{(k+1)}}} \mu_{i,0\to1}^{1-\delta_{s_i^{(k)},s_i^{(k+1)}}} \right]^{1-s_i^{(k)}} \times$$
$$\left[ (1-\mu_{i,1\to0})^{\delta_{s_i^{(k)},s_i^{(k+1)}}} \mu_{i,1\to0}^{1-\delta_{s_i^{(k)},s_i^{(k+1)}}} \right]^{s_i^{(k)}}. \tag{3}$$

Without going through the derivation in the main text, the EMF equations for $\tilde{h}_i^{(k+1)}$ remain unchanged from the main text (which do not explicitly depend on the mutation rates), while the frequencies of mutations become

$$m_i^{(k+1)} = \frac{e^{\tilde{h}_i^{(k+1)}}[\mu_{i,0\to1}(1-m_i^{(k)}) + (1-\mu_{i,1\to0}) m_i^{(k)}]}{\tilde{Z}_i^{(k+1)}\left(m_i^{(k)}; \tilde{h}_i^{(k+1)}\right)}, \tag{4}$$

(cf. Eqs (1) and (2)).

#### Inferring the mutation rate matrix for patient CH58

To apply the above method to the dynamics of p24 infecting patient CH58, we need to construct a mutation rate matrix containing $\mu_{i,0\to1}$ and $\mu_{i,1\to0}$ for $i = 1, \cdots, L$. We used the nucleotide transition matrix estimated in Zanini et al. [2] and the initial p24 nucleotide sequence of patient CH58. For sites $i$ with initial state 0, we found $\mu_{i,0\to1}$ to be the sum of nucleotide-to-nucleotide transition rates (in day$^{-1}$) that caused a change to a non-consensus amino acid. The value of $\mu_{i,1\to0}$ is small in principle because if a codon is Hamming distance 1 away from consensus, the majority of additional mutations would take it further away. We thus set $\mu_{i,1\to0}$ to be a small number $10^{-15}$ (which is of order $\mu^3$).

For sites $i$ with initial state 1, we found $\mu_{i,1\to0}$ to be the combined rate of all single nucleotide substitutions (or all double substitutions if one is not possible, or all triple substitutions otherwise) leading to the amino acid mutating to the site $i$ consensus. Similar to the above, we set $\mu_{i,0\to1} = 10^{-15}$ for these sites.

#### Results

Figure 1 shows the dynamics at sites 15, 110 and 116 within epitopes, and at sites outside epitopes (with selected sites labeled). The biggest difference we found was that reversions occur more slowly. In particular, because two mutations were required to mutate the initial codon of patient CH58 at site 91 to the consensus amino acid, it was not observed to revert during the timescale simulated. While the dynamics obtained here are similar to the constant-$\mu$ case considered in the main text, we do not expect it to be true in general: for example, escape mutations requiring rare transitions would take longer to emerge.

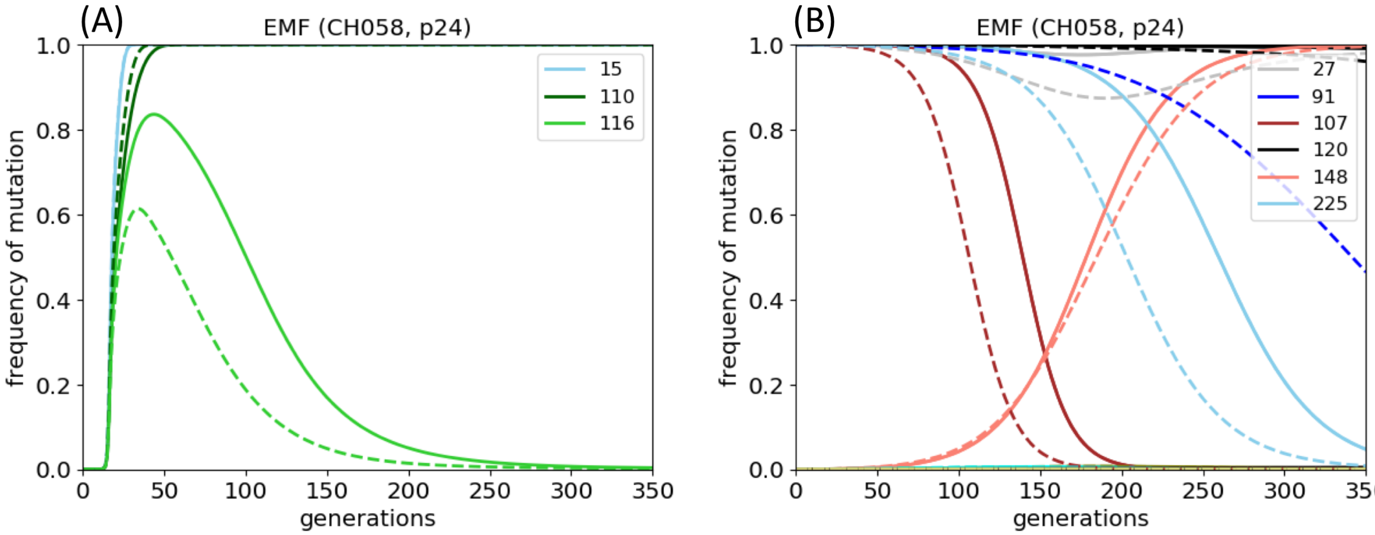


**Fig 1. Frequencies of mutations for EMF with state and site-dependent mutation rates.** Frequencies of mutations at (A) sites 15, 110 and 116 within epitopes, and (B) sites outside epitopes, as predicted by the EMF method with state and site-dependent mutation rates (solid lines), and with a uniform mutation rate $\mu^* = 3.6 \times 10^{-5}$ day$^{-1}$ as in the main text (dashed lines). Note that the solid and dashed lines for site 15 are overlapping. The within-epitope dynamics are very similar, but reversions occur more slowly. In particular, site 91 does not revert during the simulated timescale because $\mu_{91,1\to0} \ll \mu^*$ (since two mutations were required to mutate the initial codon of patient CH58 to the consensus amino acid at that site).
