## Supplementary material for "Mean-field computational approach to HIV dynamics on a fitness landscape": S2 Appendix

### Validation of EMF dynamics results by Wright–Fisher simulations

In the main text, we used the EMF method to compute the dynamics of the HIV p24 protein infecting host CH58. Barton et al. [1] computed these dynamics using fixed-population-size stochastic (Wright–Fisher) simulations, and found a good statistical correlation between the locations and relative rates at which escape mutations arise *in silico* and observed in patients (taken from Liu et al. [2]). Stochastic simulations require performing many runs before one can start making predictions of the locations and timescales of HIV escape mutations, and after the fact it is unclear why certain mutations occur and not others. EMF overcomes these issues as it is a quick semi-analytical computation involving iterating effective fitnesses and frequencies of mutations forward in time, and mutations likely to fix are directly encoded in the effective fitnesses, as explained in the main text. Here, we validate that the dynamics we obtained via EMF also arise in Wright–Fisher simulations for the example that we study.

Following Barton et al. [1], we consider a fixed population size of $N = 10^4$. During each generation, each sequence in the population has a number of offspring proportional to $e^{F(S)}$, and among all of the offspring, $N$ individuals are resampled to form the population at the next generation. (Barton et al. specify a slightly different functional form for the probability of survival of a sequence in the next generation [1].) In each generation, each offspring sequence has mutations with probability $\mu$/site, and sequences recombine at a rate $\rho$/site (we set $\mu = 3.6 \times 10^{-5}$/site/generation and $\rho = 1.4 \times 10^{-5}$/site/generation [3]).

#### Results

The mean frequencies of mutation within each epitope, over 500 simulation runs, are shown in Fig 1 (cf. Fig 7 of the main text), showing that escape mutations at sites 15 and 110 also arise. Also, Fig 2 shows what happens when $b_{1,0} = b_{2,0}/5$ and when the infecting sequence is the NL4-3 strain (cf. Fig 8 of the main text). Immunodominance indeed changes the order at which the mutations arise in the two epitopes, and infecting with the NL4-3 strain again caused site 116 instead of site 110 to mutate. However, in all of these cases, the dynamics occur more slowly than for EMF, which is to be expected because EMF describes the high-recombination-rate limit.

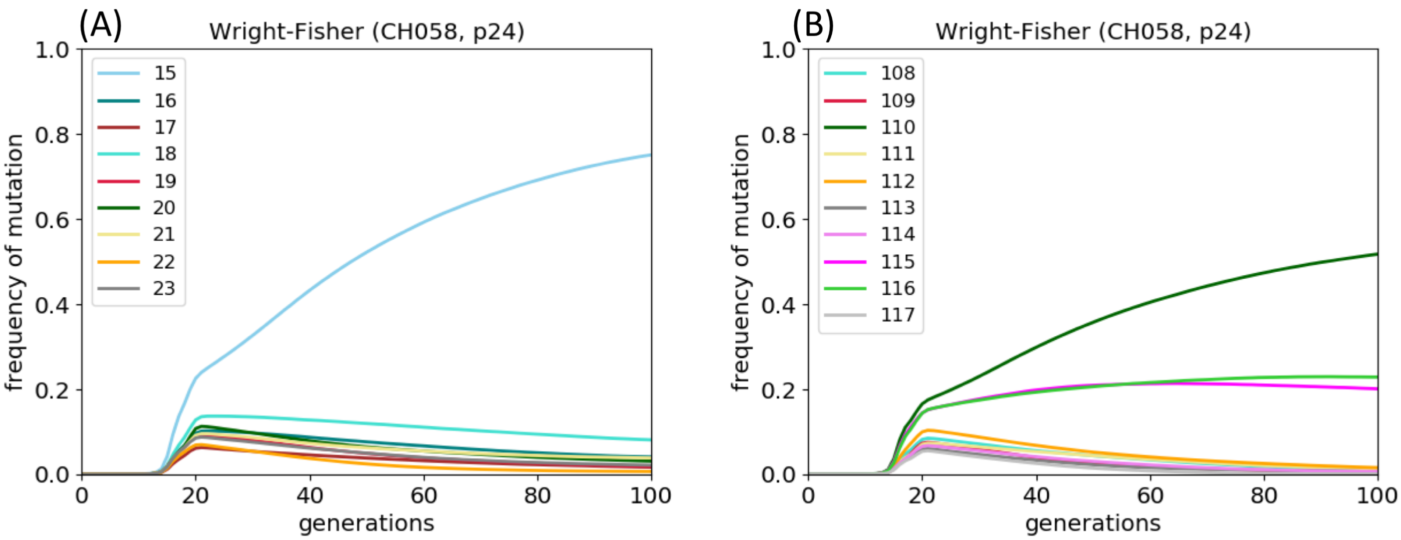


**Fig 1. Frequencies of mutations within p24 epitopes targeted by patient CH58 at sites (A) 15–23, and (B) 108–117, as predicted by Wright–Fisher simulations.**

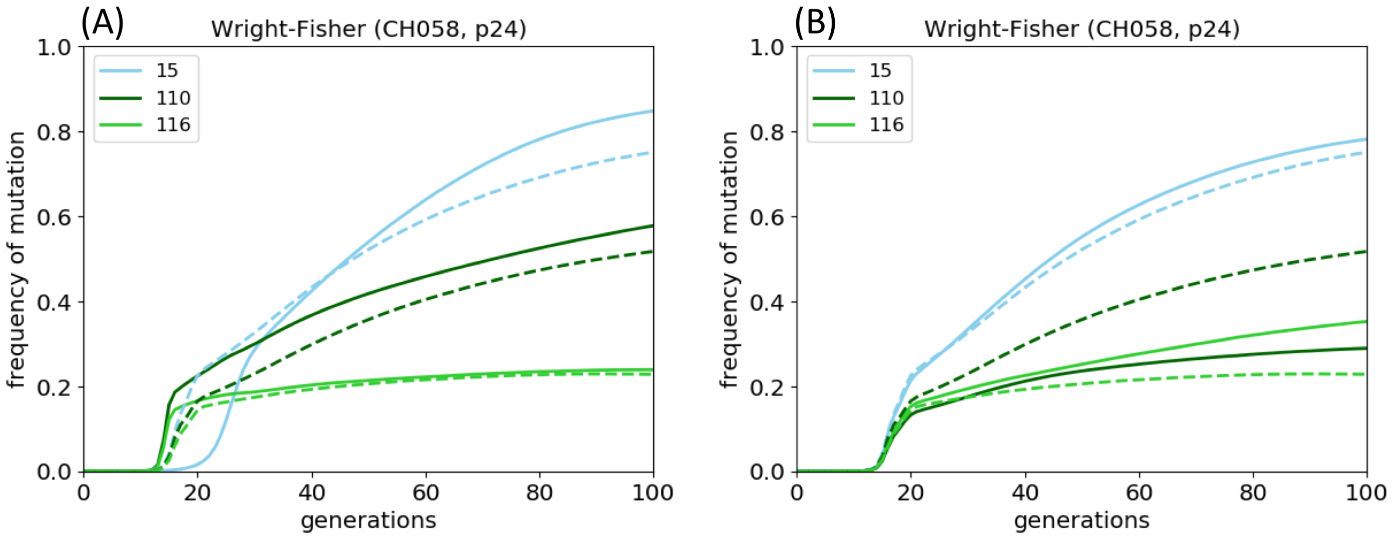


**Fig 2. Effect of immunodominance and sequence background on HIV dynamics within patient CH58, as predicted by Wright–Fisher simulations.** (A) Mutational dynamics at sites 15, 110 and 116 when $b_{1,0} = b_{2,0}/5$ (solid lines) and when $b_{1,0} = b_{2,0}$ (dotted lines), for the same $b_{\text{tot}} = b_{1,0} + b_{2,0}$. (B) Mutational dynamics at sites 15, 110 and 116 when the founder sequence is the NL4-3 strain ($s_{120}$ and $s_{208} = 1$). Site 116 mutates instead of 110 according to Wright–Fisher simulations, agreeing with the predictions from EMF.
