## Supplementary material for "Mean-field computational approach to HIV dynamics on a fitness landscape": S1 Fig

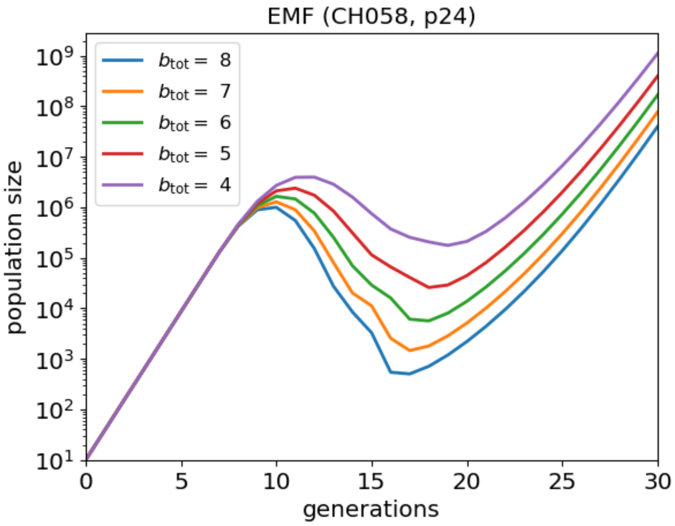


**Fig S1. EMF population dynamics produces a characteristic exponential rise and fall of population size during the early stages of infection, for a range of $b_{\text{tot}}$.**
